# Androglobin, a chimeric mammalian globin, is required for male fertility

**DOI:** 10.1101/2021.09.16.460596

**Authors:** Anna Keppner, Miguel Correia, Sara Santambrogio, Teng Wei Koay, Darko Maric, Carina Osterhof, Denise V Winter, Angèle Clerc, Michael Stumpe, Frédéric Chalmel, Sylvia Dewilde, Alex Odermatt, Dieter Kressler, Thomas Hankeln, Roland H. Wenger, David Hoogewijs

## Abstract

Spermatogenesis is a highly specialised process, involving multiple dedicated pathways and regulatory check-points. Defects ultimately lead to male sub-fertility or sterility, and numerous aspects of mammalian sperm formation remain unknown. The predominant expression of the latest globin family member, androglobin (Adgb) in mammalian testis tissue prompted us to assess its physiological function in spermatogenesis. Adgb knockout mice display male infertility, reduced testis weight, impaired maturation of elongating spermatids, abnormal sperm shape and ultrastructural defects in microtubule and mitochondrial organisation. Epididymal sperm from Adgb knockout animals display multiple flagellar malformations including coiled, bifide or shortened flagella, and erratic acrosomal development. Following immunoprecipitation and mass spectrometry, we could identify septin 10 (Sept10) as interactor of Adgb. The Sept10-Adgb interaction was confirmed both *in vivo* using testis lysates, and *in vitro* by reciprocal co-immunoprecipitation experiments. Furthermore, absence of Adgb leads to mislocalisation of Sept10 in sperm, indicating defective manchette and sperm annulus formation. Finally, *in vitro* data suggest that Adgb contributes to Sept10 proteolysis in a calmodulin (CaM)-dependent manner. Collectively, our results provide evidence that Adgb is essential for murine spermatogenesis and further suggest that interdependence between Adgb and Sept10 is required for sperm head shaping via the manchette and proper flagellum formation.

## Introduction

Spermatogenesis is a complex and tightly regulated process, involving germ cell differentiation, haploid cell formation and sperm maturation and elongation (Hermo et al. 2010a, b, c, d, e, Neto et al. 2016). In a first proliferative step, germ cells will either self-renew to maintain the pool of active progenitor cells, or commit to become spermatogonia. Spermatogonia then undergo mitosis, thereby generating primary spermatocytes, that will go through a first round of meiosis, forming secondary spermatocytes, and a second round of meiosis to generate spermatids (Mecklenburg and Hermann 2016). At this stage, DNA replication is arrested and all the genetic material (RNA) required for spermatid maturation and further development is stored in a specialized compartment known as the chromatoid body (Peruquetti 2015). During this maturation process, known as spermiogenesis, the spermatids undergo deep morphological changes, including the formation of the acrosome (the sperm head) from the Golgi apparatus before migration of the latter to the cytoplasmic droplet (Khawar, Gao, and Li 2019), condensation of the nucleus, and the sperm flagellum formation from the centriole, around which mitochondria will migrate to form the midpiece, the sperm annulus and the mobile tail (Hermo et al. 2010a, b, c, d, e). These different steps ultimately lead to male fertility and are governed by numerous factors, including hormonal, translational, post-translational and epigenetic events. Dysfunctions in any of these regulatory processes will have consequences on sperm quality, sperm motility, or fecundation capacity (Neto et al. 2016). However, numerous physiological and pathological aspects of spermatogenesis are still not fully understood, and about 30% of sperm abnormalities are still idiopathic (Fainberg and Kashanian 2019). Moreover, several studies suggest that semen quality and male fertility have declined in Western countries over the past 50 years, further urging the need of a better global understanding of male gamete formation (Levine et al. 2017). This led in recent years to the identification of a growing number of genes implicated in spermatogenesis and appearing causative for partial or complete male infertility upon mutation (Bracke et al. 2018a).

Globins are small globular metallo-proteins, which have the capacity to reversibly bind gaseous ligands via a typical 8 alpha-helical structure in which a heme prosthetic group can be embedded. In mammals, five globin types exist: the well-established hemoglobin (Hb) and myoglobin (Mb), neuroglobin (Ngb) in neuronal cells, cytoglobin (Cygb) ubiquitously expressed in fibroblasts, and the more recently identified androglobin (Adgb), predominantly expressed in mammalian testis tissue (Keppner et al. 2020).

Adgb is a chimeric protein containing an N-terminal calpain-like cysteine protease domain, a central permuted functional globin domain (Bracke, Hoogewijs, and Dewilde 2018b), interrupted by a potential calmodulin (CaM)-binding IQ motif and a large 700 amino acid long C-terminal tail of unknown identity (Hoogewijs et al. 2012). Decreased mRNA expression levels in infertile versus fertile men (Platts et al. 2007) suggest a potential role of Adgb in spermatogenesis. Gene regulation and expression studies further suggest an association of Adgb with ciliogenesis including flagellum formation (Koay et al. 2021). However, the *in vivo* function of Adgb remains unexplored. In this study we investigated the physiological function of Adgb during murine spermatogenesis by generating and analyzing Adgb knockout mice. We show that Adgb is mainly expressed in late steps of spermiogenesis, that it locates to the acrosome, the sperm flagellum, the annulus and the midpiece, and that it is crucial for male fertility and sperm formation. Furthermore, we demonstrate that Adgb interacts with septin10 (Sept10), and that co-localization occurs from the first steps of acrosome formation onwards within the sperm neck and annulus. Finally, *in vitro* data suggest that Adgb contributes to Sept10 proteolysis in a CaM-dependent manner.

## Results

### Adgb knockout mice display male infertility

A gene-trap strategy, provided by the Knockout Mouse Project (KOMP)(Skarnes et al. 2011), was applied to target exons 13 and 14 of the Adgb gene (**Fig S1**). The correct targeting of ES cells was verified first by long-range PCR (**Fig. S1**), and second by Southern blotting (**Fig. S1**). The targeted Adgb^tm1a(KOMP)Wtsi^ allele (Adgb tm1a mice), generated on a C57BL/6N background, displays a gene-trap DNA cassette, which was inserted into the twelfth intron of the *Adgb* gene. The gene trap consists of a splice acceptor site, an internal ribosome entry site, a β-galactosidase reporter sequence, and a neomycin resistance sequence. Breeding of Adgb tm1a mice with ubiquitously expressed CMV Cre-deleter mice allowed generation of mice deficient for exons 13 and 14 but still expressing the β-galactosidase reporter (Adgb tm1b mice) (**Fig. S1**). Furthermore, mating of Adgb tm1a mice with Flp-deleter mice enabled the generation of conditional floxed mice (Adgb tm1c) (**Fig. S1**). These mice were further crossed with CMV Cre-deleter mice to generate the full knockout animals (Adgb tm1d) (**Fig. S1**), that were used for all downstream applications if not otherwise stated. Genotyping was performed by regular PCR (**Fig. S1**), and revealed no significant differences in the Mendelian distribution of offspring for Adgb tm1d animals following interbreeding of heterozygous parents (380 pups: +/+, n= 101; +/-, n= 168; -/-, n=111; χ^2^ = 5.62, p > 0.05, ns). The genetic ablation of Adgb expression was further verified by RT-qPCR (**Fig. 1A**) and immunoblotting (**Fig. 1B,C**). While female knockout mice displayed no fertility issues, male knockout mice never generated offspring, indicative for infertility. Full penetrance male infertility was also observed in homozygous tm1a and tm1b male mice, whereas homozygous tm1c animals showed normal fertility (data not shown). Accordingly, the testis weight was significantly reduced in knockouts (**Fig. 1D**) associated with decreased serum testosterone levels (**Fig. 1E**). Stage-specific histological examination of seminiferous tubules of control animals revealed normal architecture, normal spermatogenic maturation steps, and the presence of mature sperm with flagella extending into the lumen (**Fig. 1F**). In contrast, in knockout animals, despite the presence of meiotic events (stages X-XII, **Fig. 1F**), no flagella could be observed during the spermiation stage (stage VII-VIII, **Fig. 1F**). The absence of mature sperm was accompanied by abnormally shaped heads, trapped stage 16 spermatids within the epithelium and the presence of cytoplasmic material filling the lumen of the tubules (**Fig. 1F**). Within the cauda epididymis, knockout animals displayed accumulations of residual bodies, cytoplasmic material, shed germ cells and occasional abnormally shaped sperm heads, but an overall absence of mature sperm as compared to wildtype animals (**Fig. 1G**). No differences were detected in knockout testes at mRNA levels of nitric oxide synthases 1-3 (NOS1-3) or superoxide dismutases 1-3 (SOD1-3), and the ratio of Bax/Bcl2 was unchanged compared to wildtype testes, suggesting no increase in apoptotic events or oxidative stress (**Fig. S2**).

**Figure 1.**
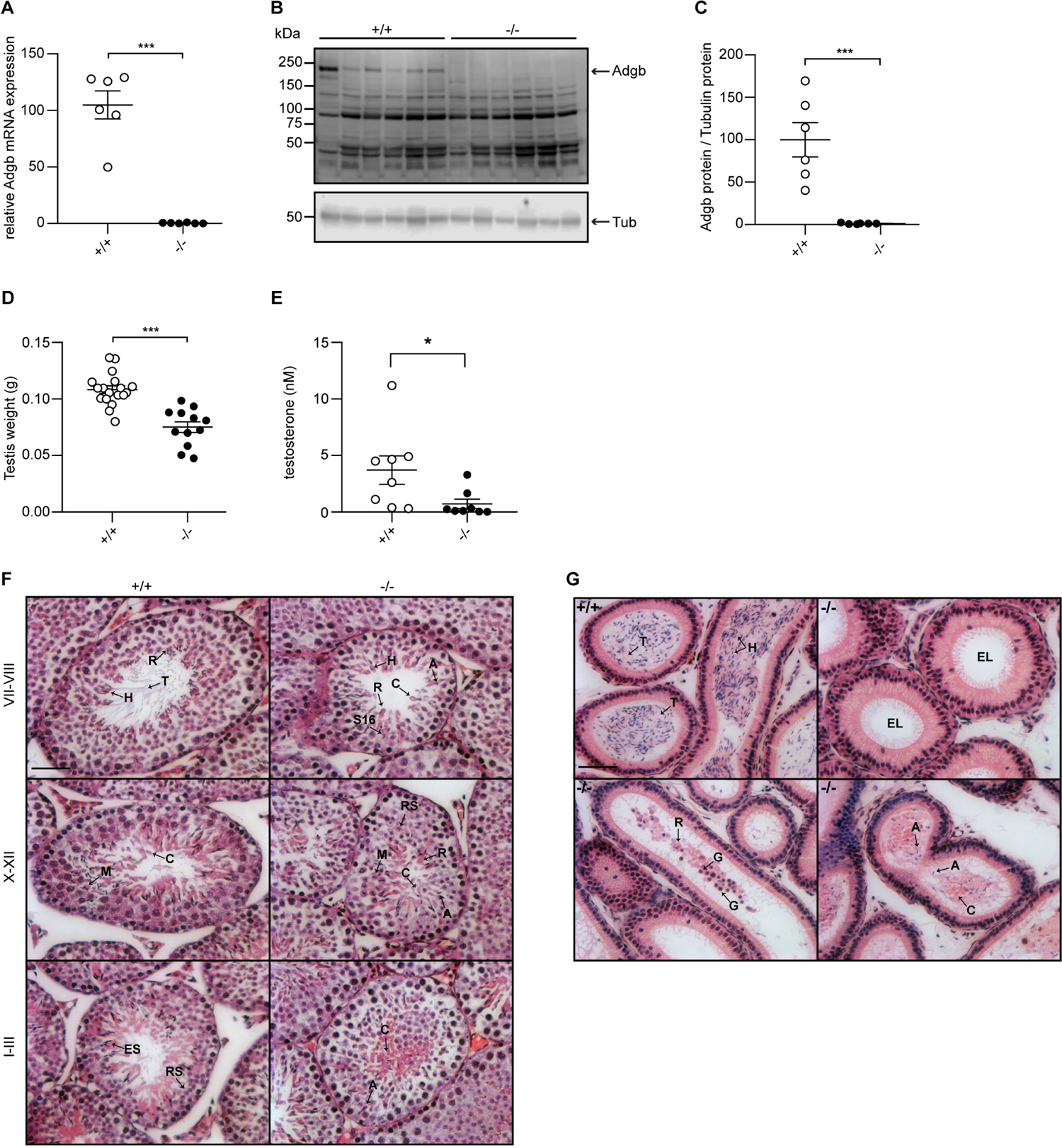
Validation of the knockout model and testicular phenotype. (**A**) Relative mRNA expression levels of Adgb in testes of wildtype (+/+) and knockout mice (-/-) (n=6 per genotype; p=0.000008). (**B**) Representative immunoblot for Adgb in testis lysates from wildtype (+/+) and knockout mice (-/-) (n=6 per genotype) and (**C**) corresponding protein quantification. Tubulin was used as loading control. p=0.0007. (**D**) Testis weight (g) in Adgb wildtype (+/+), heterozygous (+/-) and knockout (-/-) mice (n=8-13 per genotype). p=0.000003. (**E**) Serum testosterone levels (nM) in Adgb wildtype (+/+) and knockout (-/-) mice (n=8 per genotype). p=0.041. (**F**) Representative hematoxylin and eosin (H&E) stained sections of testes from Adgb wildtype (+/+) and knockout mice (-/-) at the different stages of spermatogenesis. Heads (H), tails (T), residual bodies (R) cytoplasmic bulges (C), meiosis (M), elongating spermatids (ES), round spermatids (RS), stage 16 spermatids (S16) and abnormal heads (A) are indicated. Scale bar represents 50 µm. (**G**) Representative H&E stained sections of epididymides from Adgb wildtype (+/+) and knockout mice (-/-). Heads (H), tails (T), cytoplasmic bodies (C), residual bodies (R), germ cells (G), abnormal heads (A) are shown. Note the empty lumen (EL) in knockout mice. Scale bar represents 50 µm. ** p< 0.01, *** p< 0.001.

### Absence of Adgb interferes with the maturation of elongating spermatids

To determine the temporal expression pattern of Adgb during spermatogenesis, wildtype embryos and pups at different post-natal ages, corresponding to the stages of the first wave of spermatogenesis during puberty, were dissected and analysed by RT-qPCR. Whereas the expression of Adgb nearly remained undetectable until post-natal day 21 (corresponding to the stage of round spermatids), Adgb mRNA levels drastically increased to reach a peak at post-natal day 25, coinciding with the first elongating spermatids (**Fig. 2A**). Bulk and single-cell RNA sequencing analysis in mouse and human datasets available at the ReproGenomics Viewer resource (Darde et al. 2015, Darde et al. 2019) confirmed the conserved expression pattern of ADGB in spermatids (Green et al. 2018, Jégou et al. 2017, Lukassen et al. 2018, Wang et al. 2018) (**Fig. S3**). Accordingly, Adgb protein expression equally reached its peak at post-natal days 26 to 28 (**Fig. 2B,C**). This finding was further confirmed by propidium iodide staining and FACS sorting on testis lysates of the different genotypes. While no variations could be detected for phases 2C (spermatogonia, secondary spermatocytes, testicular somatic cells), S-phase (pre-meiotic spermatogonia), 4C (primary spermatocytes) and 1C (round spermatids), an abnormal accumulation of elongating and elongated spermatids (phase H) could be detected in knockout animals, suggesting a blockade in the elongation process (**Fig. 2D**). Additionally, immunofluorescence (**Fig. 2E**), mRNA *in situ* hybridization (**Fig. 2F**) and X-gal (**Fig. 2G**) stainings of testis sections from wildtype and knockout mice confirmed the presence of Adgb within layers containing post-meiotic cells, and further intensifying towards the lumen and mature sperm (**Fig. 2E-G**). Moreover, in mature sperm, Adgb expression could be visualized within the midpiece and along the whole flagellum by both X-gal (**Fig. 2F**) and immunofluorescence (**Fig. 2G**) stainings.

**Figure 2.**
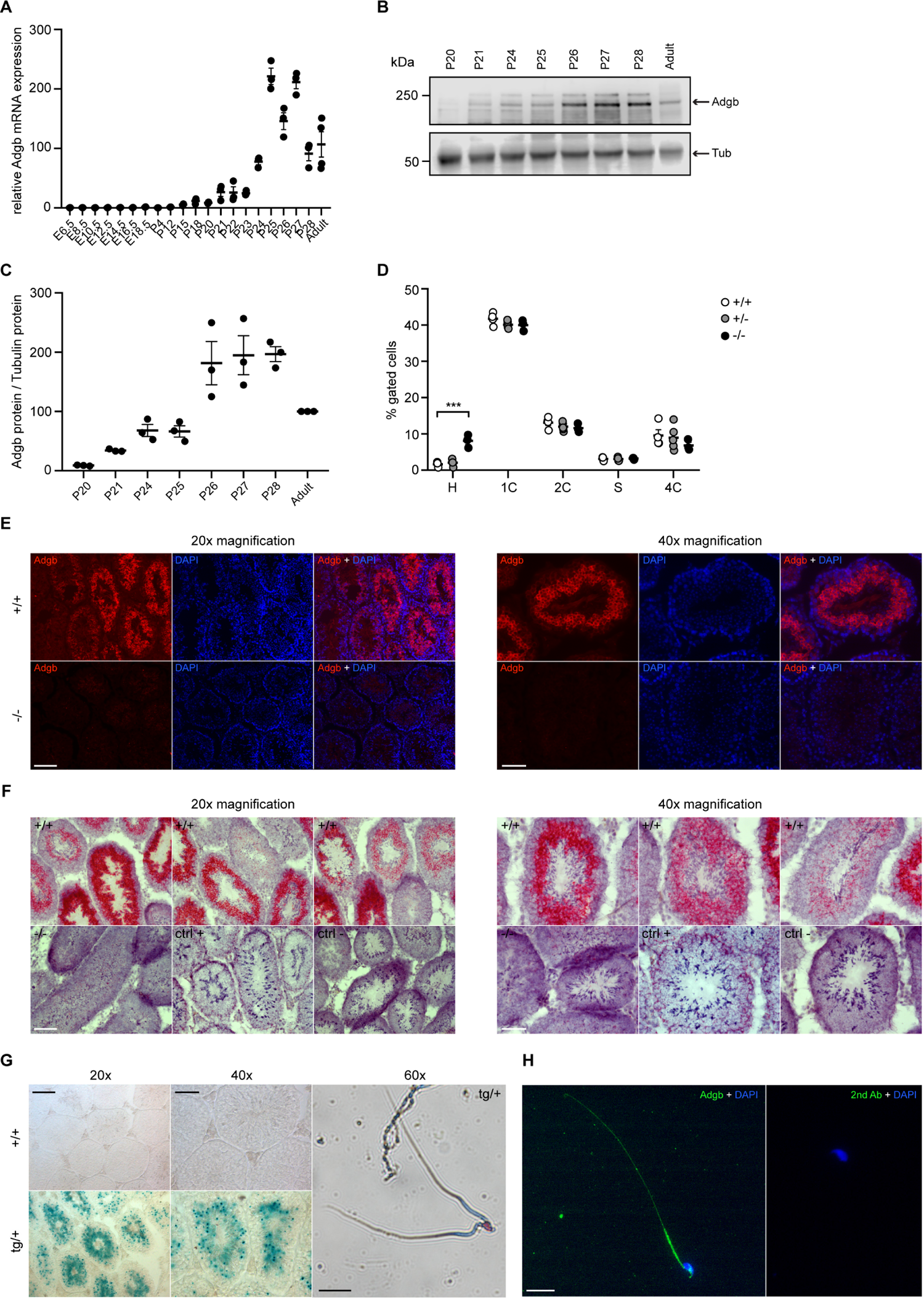
Testicular Adgb expression pattern and localization. (**A**) Relative mRNA expression levels of Adgb in testes of wildtype mice during embryonic development (E) and early post-natal (P) life (n=3-4 per condition). (**B**) Representative immunoblot for Adgb in testis lysates from wildtype mice at different post-natal (P) ages (n=3 per condition) and (**C**) corresponding protein quantification. Tubulin was used as loading control. (**D**) Flow cytometric analysis of spermatogenic cell populations following propidium iodide staining in Adgb wildtype (+/+, white circles, n=4), heterozygous (+/-, grey circles, n=5) and knockout (-/-, black circles, n=3) testes. H: elongating and elongated spermatids; 1C: round spermatids; 2C: spermatogonia, secondary spermatocytes, testicular somatic cells; S: spermatogonia synthesizing DNA; 4C: primary spermatocytes. p=0.00024. (**E**) Representative pictures of Adgb protein (red fluorescence) detection in testes of wildtype (+/+) and knockout (-/-) animals. Left panels 20x magnification, right panels 40x magnification, scale bars represent 100 µm and 50 µm, respectively. Nuclei were stained with DAPI. (**F**) Representative pictures of Adgb mRNA *in situ* hybridization in testes from wildtype (+/+) and knockout (-/-) animals. Left panels 20x magnification; right panels, 40x magnification; scale bars represent 100 µm and 50 µm, respectively. Positive (ctrl +, PPIB) and negative (ctrl -, DapB) control sections are shown. (**G**) Representative pictures of b-galactosidase activity (X-gal staining) in testes from Tm1b wildtype (+/+) and Tm1b heterozygous (tg/+) mice, and isolated spermatozoa from Tm1b heterozygous (tg/+) mice. Left panels, 20x magnification; middle panels, 40x magnification; right panel, 60x magnification; scale bars represent 100 µm, 50 µm and 20 µm, respectively. Spermatozoa were counterstained with nuclear fast red. (**H**) Representative picture of Adgb protein (green fluorescence) in a single spermatozoon from wildtype (+/+) mice (left panel), and negative control (secondary antibody only, right panel). Scale bar represents 20 µm, nuclei were stained with DAPI. ** p< 0.001.

*Adgb is required for proper assembly of microtubules and positioning of mitochondria* To gain additional insights into the origin of male infertility, cauda epididymal sperm was collected from both wildtype and knockout mice, and visualized under a microscope. While wildtype sperm appeared normal, very few knockout spermatozoa were found and displayed various defects of the head and/or flagellum structure, including shortened or bifide tails, loopings of the flagellum, and immature acrosomal structures (**Fig. 3A**). Knockout sperm acrosome structure appeared partially conserved, with sperm displaying either normal acrosomes, or various defects in shape or staining intensity, or total absence (**Fig. 3B**). TEM ultrastructural analysis of testis sections further revealed misshaped heads with nuclear inclusions, disorganised axonemes failing to arrange around the flagella, misaligned microtubules, defective manchette elongation, and chaotic mitochondria within the forming midpieces of elongating spermatids in knockout samples (**Fig. 3C**).

**Figure 3.**
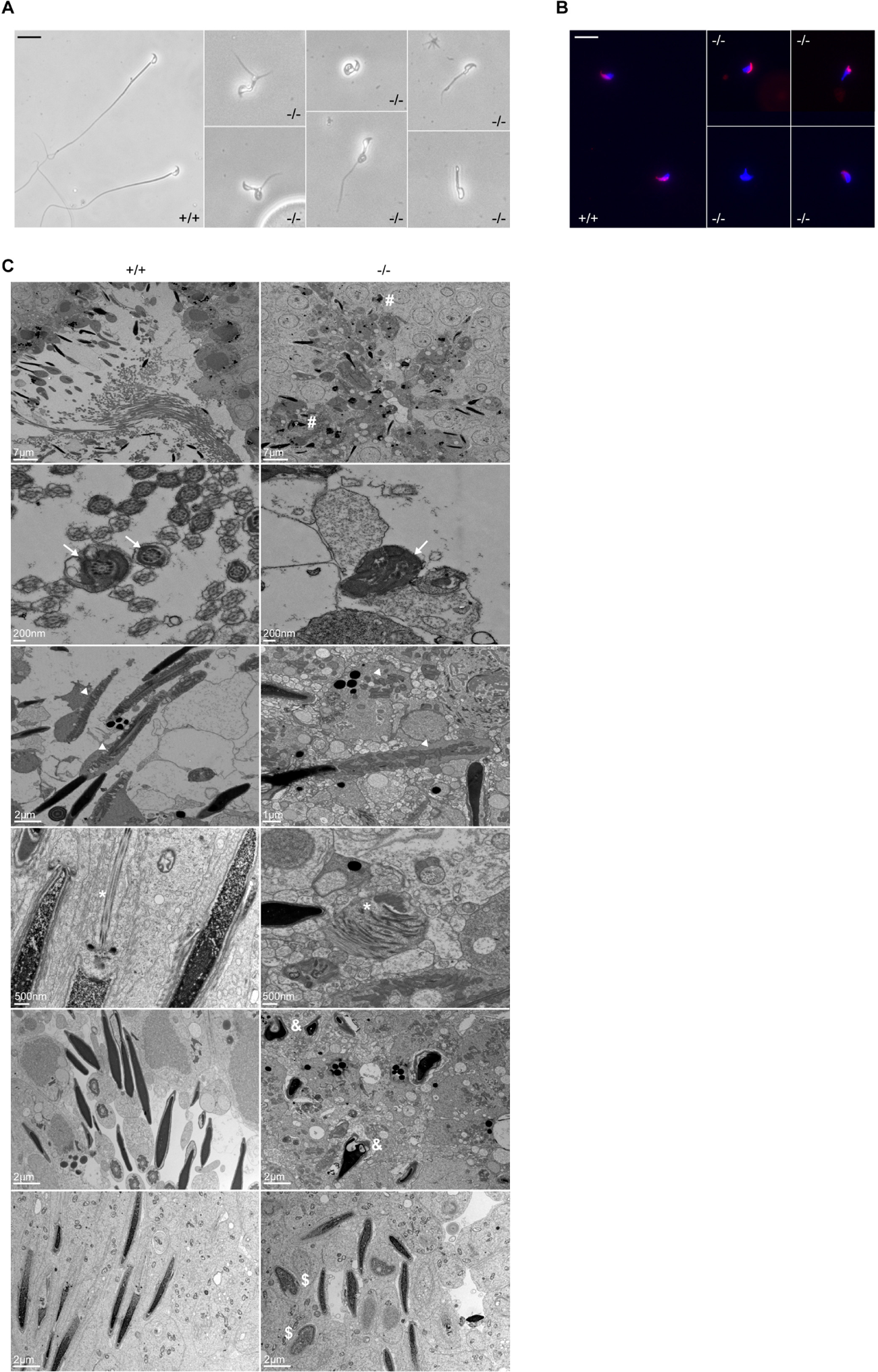
Defective spermatogenesis is associated with flagellar malformation in Adgb knockout mice. (**A**) Representative pictures of cauda epididymis sperm from wildtype (+/+) and Adgb knockout animals (-/-). Scale bar represents 20 µm. (**B**) Representative pictures of PNA-stained cauda epididymis sperm from wildtype (+/+) and Adgb knockout animals (-/-). Nuclei were stained with DAPI. Scale bar represents 20 µm. (**C**) Representative TEM pictures from wildtype (+/+, left panels) and knockout (-/-, right panels) testes. Misshaped sperm heads (hash), axonemes (arrows), mitochondria (arrowheads) and microtubules (asterisks), nuclear inclusions (ampersand), and abnormal manchette elongation (dollar) are shown. Scale bar lengths are indicated on each picture.

### The Adgb-dependent transcriptome reveals dysregulation of multiple spermiogenesis genes

To understand the molecular consequences of loss of Adgb in the testis, we performed RNA-sequencing experiments on total testis RNAs from wildtype and knockout mice at post-natal day 25. An elaborate set of significant differentially expressed genes (74 genes upregulated and 204 downregulated) was identified, underscoring the crucial function of Adgb in spermatogenesis (**Fig S4**, **Dataset S1**). Functional analysis based on Gene Ontology term enrichments confirmed that many of these genes are related to sperm head, acrosome reaction, acrosomal membrane, sperm motility, spermatid development and spermatid differentiation in line with the pronounced structural changes in spermatids during spermiogenesis. Intriguingly, a more refined Ingenuity pathway-based analysis (IPA) of the differentially expressed gene set provided a link to testosterone synthesis as deregulated pathway, as evidenced by reduced 17βhsd3 and Lhcgr mRNA levels in knockout mice and consistent with the observed decrease in serum testosterone levels (**Fig. 1E**, **Fig. S4**).

### Adgb interacts and co-localizes with Sept10

To obtain more insights into the physiological function of Adgb, we explored the Adgb-dependent interactome. Total protein extracts from wildtype testes were immunoprecipitated (IP) with anti-Adgb antibody or IgG and subsequently submitted to mass spectrometry (MS) analysis to reveal potential interacting proteins of Adgb.

Among the specifically enriched proteins, there were various members of the septin family of proteins, such as Sep10, Sept11, Sept2, and Sept7 (**Fig. 4A and Dataset S2**). Particular focus was put on Sept10 for further downstream experiments given its strong enrichment combined with high abundance in the immunoprecipitation. To confirm the interaction between Adgb and Sept10, reciprocal co-immunoprecipitation (co-IP) experiments were performed (**Fig. 4B,C**) on tissue extracts from wildtype and knockout testes (**Fig. 4B**) and in HEK293 cells overexpressing full-length ADGB (**Fig. 4C**) and Sept10. The results demonstrate that Adgb and Sept10 interact both *in vivo* (**Fig. 4B**) and *in vitro* (**Fig. 4C**), whereas in testis lysates of Adgb-deficient mice, no Sept10 precipitation was observed (**Fig. 4B**). Endogenous Sept10 protein levels were equal in testis lysates of Adgb-deficient and wildtype mice. Similar observations were made for Sept11, Sept7, and Sept2, as well as for other septins that are crucial for spermatogenesis, including Sept8, Sept9 and Sept14 (**Fig. S5**). We next investigated whether the interaction with Sept10 occurs at the N-terminal or the C-terminal portion of Adgb (**Fig. 4D,E**). Following co-overexpression of Adgb deletion constructs with SEPT10 and subsequent co-immunoprecipitation, immunoblotting revealed that both parts of ADGB interact with SEPT10 (**Fig. 4D,E**), and that this interaction remained intact also upon deletion of the coiled-coil domain of ADGB (**Fig. S6**).

**Figure 4.**
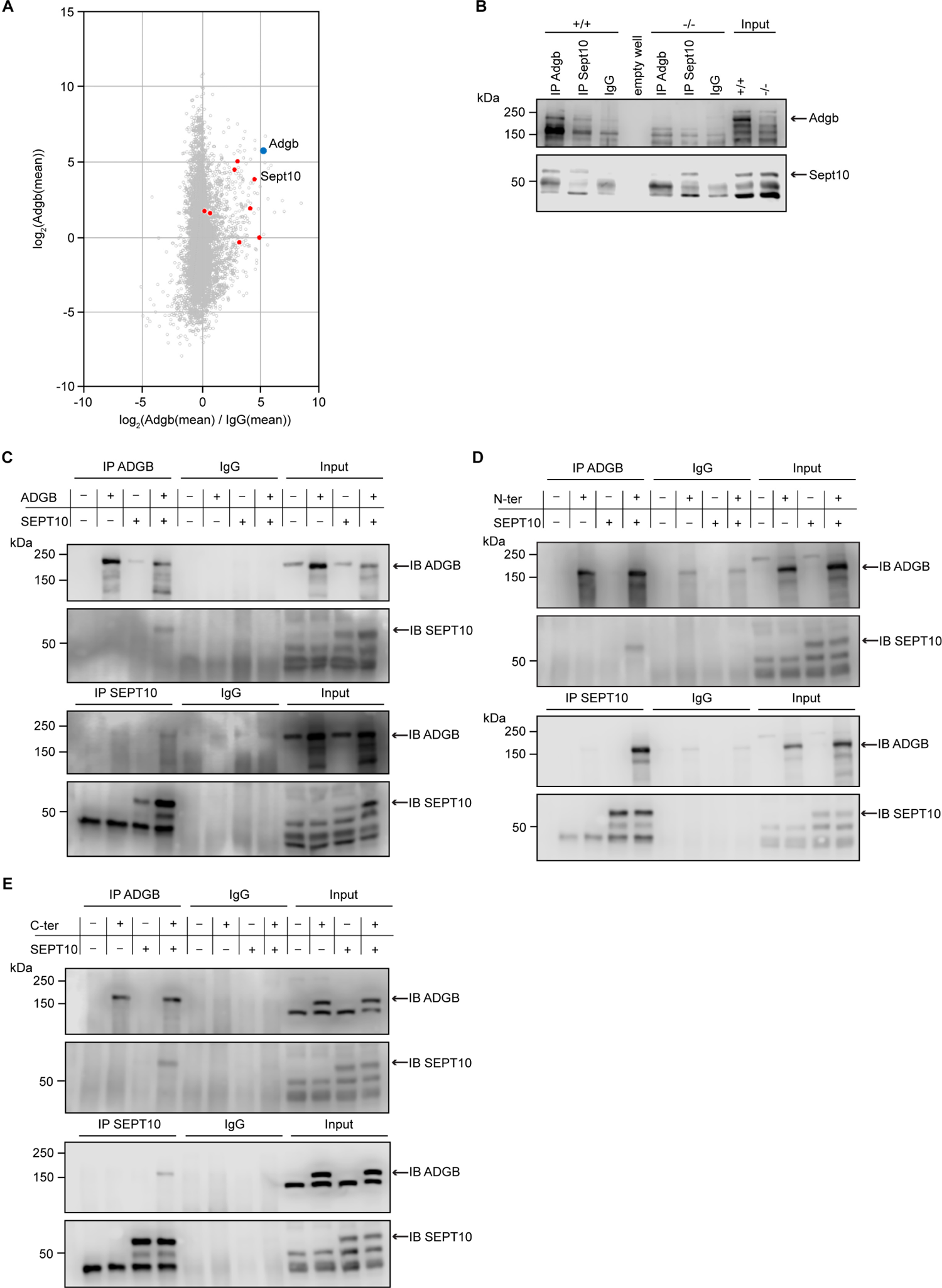
Adgb and Sept10 interact *in vivo* and *in vitro*. (A) Proteins of the septin family are specifically enriched in the Adgb immuno-precipitation (IP). The iBAQ (intensity-based absolute quantification) values of each Adgb IP (triplicate) and IgG control IP (duplicate) were log2 transformed and normalized against the median value. Missing values were imputed before the mean values of the Adgb and IgG control IPs were calculated. The normalized abundance of each protein detected in the Adgb IP (log2 Adgb (mean)) is plotted against its specific enrichment compared to the IgG control IP (log2 (Adgb (mean) / IgG (mean)). Adgb and septins are highlighted as blue and red dots, respectively, in the christmas tree plot representation. (**B**) Representative immunoblot of Adgb and Sept10 in testis lysates from wildtype (+/+) and knockout (-/-) mice following co-immunoprecipitation of Adgb and Sept10. (**C-E**) Representative immunoblots of ADGB and Sept10 in protein lysates of HEK293 cells (co-)transfected with full-length ADGB (**C**), N-ter ADGB (**D**) and C-ter ADGB (**E**) and Sept10 following co-immunoprecipitation of ADGB and Sept10.

Consistent with a functional interaction, the temporal expression profiles of SEPT10 and ADGB substantially overlapped as illustrated by analysis of bulk and single-cell RNA sequencing datasets of mouse and human RNA (**Fig. S3**) as well as by RT-qPCR and immunoblotting of mouse tissue samples (**Fig. S7**). The localization of Adgb and Sept10 was assessed in microdissected tubules and in epididymal sperm by immunofluorescence. Co-localization of Adgb and Sept10 was visible during the Golgi phase in the acrosomal granule, during manchette and sperm tail formation (**Fig. 5A**), and at the level of the sperm annulus in mature wildtype sperm (**Fig. 5B**). A moderate Sept10 staining was also observed in the neck region of mature sperm (**Fig. 5B**). In knockout epididymal sperm, only a single signal, likely corresponding to the annulus, was observed and displayed abnormal migration, indicating defective manchette or microtubule formation (**Fig. 5B**). The migration of the annulus drives mitochondrial placement along the forming mid-piece (Toure et al. 2011). Since knockout animals displayed abnormal ultrastructural mitochondria organisation (**Fig. 3C**), CoxIV staining was performed on wildtype and knockout epididymal sperm. As expected, a robust staining was observed along the whole midpiece in wildtype sperm, whereas mitochondria were barely visible and formed a cloudy structure around the neck region of knockout sperm (**Fig. 5B**).

**Figure 5.**
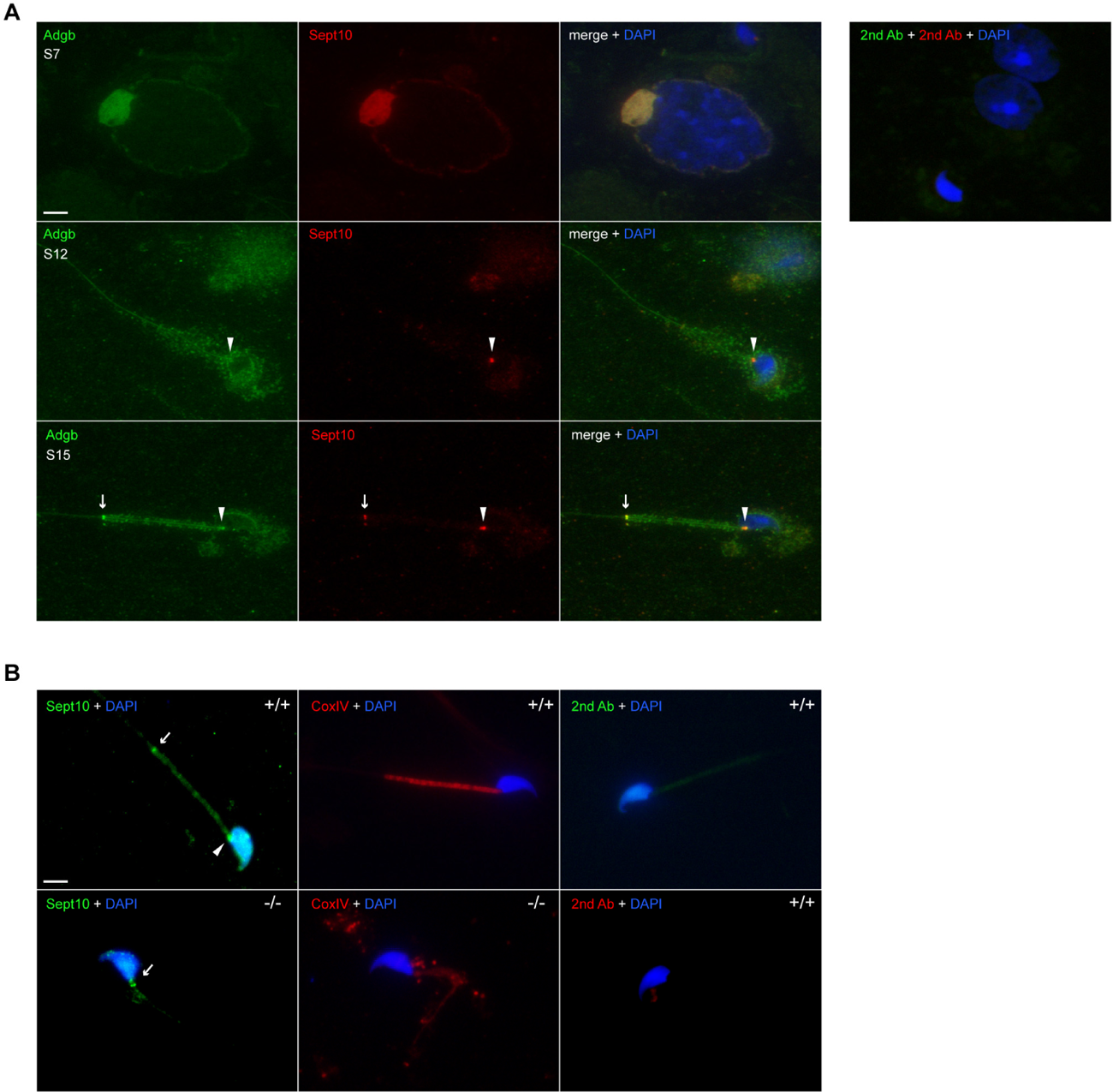
Adgb and Sept10 co-localize in developing acrosome, sperm neck and annulus. (A) Representative pictures of Adgb protein (green fluorescence) and Sept10 (red fluorescence) in elongating spermatids (stage 7 [S7] upper panels, stage 12 [S12] middle panels, and stage 15 [S15] lower panels) after stage-specific tubule dissection of wildtype testes. Nuclei were stained with DAPI. Negative control (secondary antibodies only) is shown on the right. Scale bar represents 10 µm. Sperm neck (arrowhead) and annulus (arrow) are highlighted. (**B**) Representative pictures of Sept10 (left panels, green fluorescence) and CoxIV (middle panels, red fluorescence) proteins in wildtype (+/+) and knockout (-/-) epididymal sperm. Sections were counterstained with DAPI. Negative control (secondary antibodies only) are shown (right panels). Scale bar represents 10 µm.

### ADGB contributes to Sept10 proteolytic cleavage in vitro

To analyse the functional consequences of the SEPT10-ADGB interaction we transiently co-overexpressed both proteins in HEK293 cells. Intriguingly, apart from the intact form of overexpressed SEPT10 at 60 kDa, increased levels of a lower band of 37 kDa were consistently detected in the presence of exogenous ADGB (**Fig. 6A**), in a dose-dependent manner (**Fig. 6A**). Immunoblotting with a V5-antibody upon co-overexpression of a C-terminal V5-tagged SEPT10 with ADGB (**Fig. 6B**) as well as the presence of this band upon SEPT10/ADGB co-IP (**Fig. 6B, Fig. S8**) further support its origin as proteolytic SEPT10 product. To investigate a potential influence of the globin domain this experiment was repeated under normoxic and hypoxic conditions (0.2% O_2_) but no differences were observed upon exposure to hypoxic conditions (**Fig. 6C**). To determine a potential role of calmodulin we constructed a deletion mutant lacking the IQ domain. Notably, transient overexpression of IQ-mutant ADGB resulted in considerably reduced appearance of the 37 kDa band relative to WT ADGB (**Fig. 6D**). This finding prompted us to experimentally validate the suspected CaM-ADGB interaction. Whereas co-IP experiments following overexpression of full-length ADGB did not interact with CaM under basal experimental conditions in HEK293 cells (data not shown), a truncated construct covering the globin and IQ domains displayed robust ADGB-CaM interaction (**Fig. 6E**). Consistently, MS analysis of proteins that were present in the IP of the overexpressed, isolated globin domain revealed a prominent enrichment of endogenous CaM (**Fig. S9 and Dataset S3**). Importantly, individual or double mutation of the proximal histidine (HisF8) or distal glutamine (GluE7), critical residues in the globin domain for heme coordination, did not alter the interaction, suggesting that the ADGB-CaM interaction occurs independently of heme incorporation (**Fig. 6F, Fig. S10**). As an additional layer of support for the ADGB-CaM interaction, chimeric Gal4 DNA-binding domain and VP16 transactivation domain-based fusion constructs were generated for mammalian 2-hybrid (M2H) luciferase reporter gene assays (**Fig. 6G,H**). These M2H assays performed in 2 different cell lines, HEK293 and A375, revealed up to 7.5-fold increase in luciferase activity upon co-transfection of both chimeric proteins compared to single construct transfections, providing independent evidence that ADGB interacts with CaM. To further assess the potential O_2_-dependency of the CaM-ADGB interaction, we repeated the M2H assays in A375 cells under hypoxic conditions. Exposure to hypoxic conditions (0.2% O_2_) did not alter luciferase activity, while luciferase activity of an *EPO* promoter-driven reporter gene increased, suggesting that the ADGB-CaM interaction is O_2_-independent (**Fig. 6H**). This result is consistent with the maintained ADGB-CaM interaction following mutation of critical residues in the globin domain in co-IP experiments and unchanged ADGB-dependent cleavage of SEPT10 under hypoxic conditions. Collectively, these *in vitro* data suggest a scenario in which exogenous ADGB proteolytically contributes to cleavage of exogenous SEPT10 in an O_2_-independent but CaM-dependent manner.

**Figure 6.**
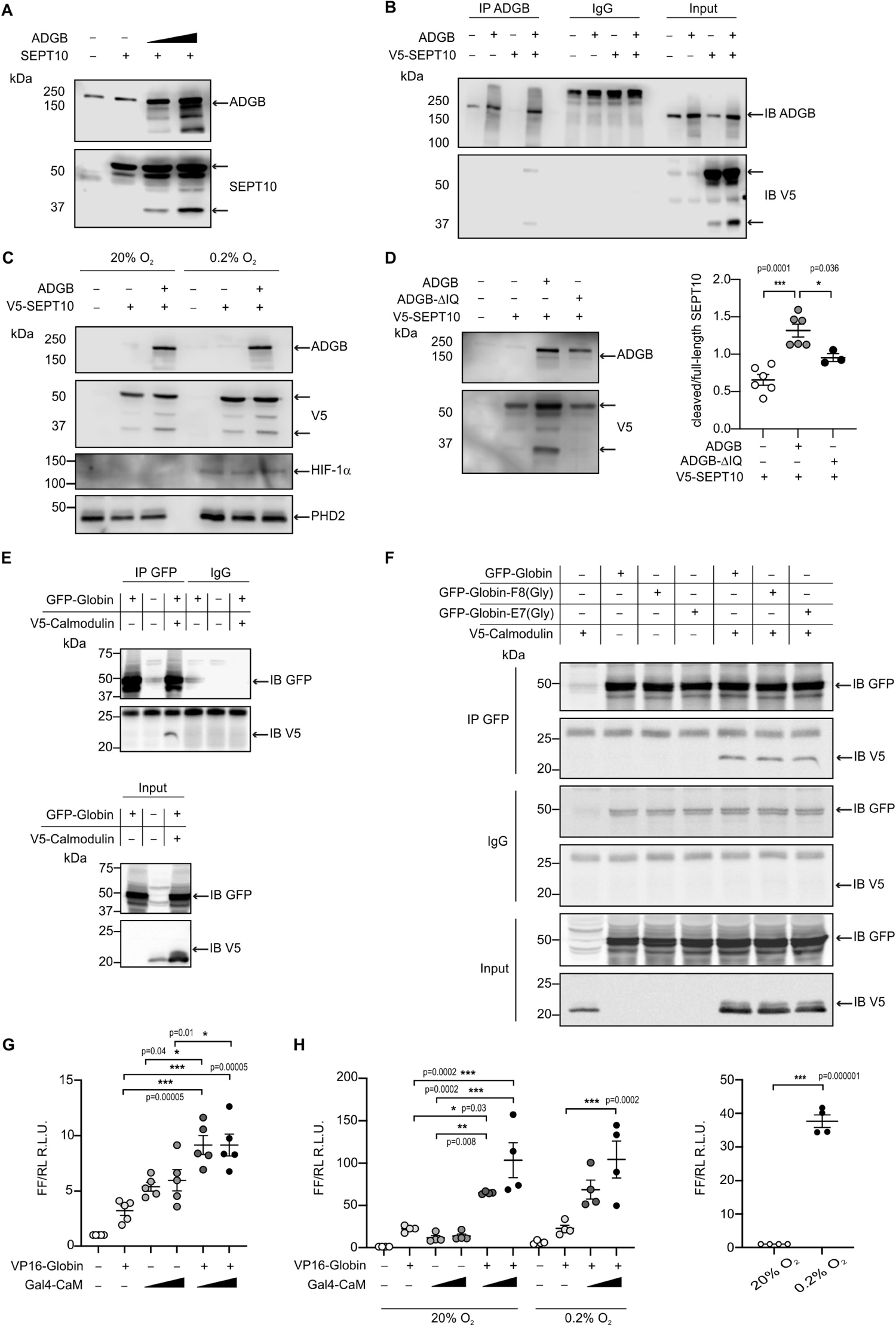
Adgb contributes to *in vitro* CaM-dependent Sept10 cleavage. **(A)** Representative immunoblots of ADGB and SEPT10 in protein lysates of HEK293 cells co-transfected with plasmids encoding SEPT10 and two dose-dependent amounts of full-length ADGB. **(B)** Representative immunoblot of ADGB and V5 in protein lysates of HEK293 cells (co-)transfected with full-length ADGB and a C-terminally V5-tagged SEPT10 construct following co-immunoprecipitation of ADGB and V5-SEPT10. (**C**) Representative immunoblots of ADGB, V5, HIF-1α and PHD2 in protein lysates of HEK293 cells (co-)transfected with full-length ADGB and V5-SEPT10 following exposure to normoxic (20% O_2_) and hypoxic conditions (0.2% O_2_) for 24 hours. HIF-1α and PHD2 were used as positive controls for hypoxia. **(D)** Representative immunoblot of ADGB and V5 in protein lysates of HEK293 cells (co-)transfected with full-length ADGB, V5-SEPT10 and an ADGB-IQ deletion mutant, and corresponding protein quantification of cleaved/full-size SEPT10 ratio (n=3-6 independent experiments). Ponceau S protein staining was used as loading control. **(E)** Representative immunoblot of GFP and V5 in protein lysates of HEK293 cells (co-)transfected with a truncated construct of the globin domain of ADGB (spanning the IQ domain) (GFP-Globin) and a V5-tagged CaM (V5-Calmodulin) following immunoprecipitation of GFP. **(F)** Representative immunoblot of GFP and V5 in protein lysates of HEK293 cells (co-)transfected with GFP-Globin, a GFP-Globin construct with mutation of the proximal heme-binding histidine (GFP-Globin-F8(Gly)), a GFP-Globin construct with mutation of the distal glutamine (GFP-Globin-E7(Gly)) and V5-CaM following immunoprecipitation of GFP. **(G, H)** Mammalian-2 hybrid assays in HEK293 cells under normoxic conditions (**G**) and A375 cells under normoxic and hypoxic (0.2% O_2_) conditions (**H**) (n=3-5 independent experiments). HEK293 and A375 cells were transiently transfected with fusion protein vectors based on a Gal4 DNA binding domain fused to calmodulin (Gal4-CaM) and a VP16 activation domain fused to the ADGB globin domain comprising the IQ domain (VP16-Globin), a Gal4 response element-driven firefly luciferase reporter, and a *Renilla* luciferase control vector. Increasing transfection amounts for the Gal4-CaM fusion protein were employed. Following transfection, A375 cells were incubated under normoxic (20% O_2_) or hypoxic (0.2% O_2_) conditions, and luciferase reporter gene activities were determined 24 hours later. Single construct transfections served as negative controls. The single Gal4-CaM control condition is not displayed in A375 cells exposed to hypoxia, due to its hypoxic regulation. An Epo hypoxia response element-driven firefly luciferase construct served as hypoxic control. * p< 0.05, ** p< 0.01, *** p< 0.001.

## Discussion

In the present study, we aimed to explore the testicular function of Adgb by generating and analysing Adgb^-/-^ constitutive knockout mice. Our results demonstrate that Adgb is required for late stages of spermatogenesis and male fertility, as evidenced by a total lack of functional sperm in knockout animals. Also, a reduction in testis weight, reduced testosterone production and dysregulated testosterone-associated genes such as Lhcgr and 17βhsd3 were observed. This finding is remarkable as Adgb displays little to no expression levels in the testosterone-producing Leydig cells. Our data further show that Adgb is indispensable for proper microtubule formation, mitochondria localisation and annulus positioning. Spermatid elongation is marked by profound morphological changes, which start with the formation of the acrosome, concomitant with the first expression of Adgb. Adgb mRNA and protein are detectable from post-natal day 21 onwards, the stage at which the acrosomal granule is formed (Clermont and Leblond 1955, Khawar, Gao, and Li 2019). Accordingly, a strong signal was visible within the acrosomal granule, and Adgb could be detected in all steps of spermatogenesis within the acrosome, including in mature sperm. Moreover, the defects in head shape and acrosome structure visible in Adgb mutant mice strongly suggest a function of Adgb in the maturation of the acrosome.

To understand the molecular mechanisms by which Adgb may exert its function during spermatogenesis we analysed the interactome of Adgb by LC-MS/MS after immunoprecipitation of the latter in testicular lysates. Strikingly, several members of the septin family ranked in the top hits. Septins are conserved GTPases that have the ability to form large oligomers and filamentous polymers and which associate with cell membranes and with the cytoskeleton. They serve as scaffolds for the proper localization of intracellular proteins via their diffusion barrier-forming characteristic. All septins display a conserved structure formed by an N-terminal proline-rich polybasic region which interacts with membrane phospholipids, a central GTP-binding domain, a C-terminal region of unknown function (the septin unique element) further flanked by a C-terminal tail containing α-helical coiled coils of varying sizes enabling the polymer-forming protein-protein interactions typical of septins (Dolat, Hu, and Spiliotis 2014).

Thirteen septins have been described so far in mammals (Sept1-12 and Sept14), further sub-classified into 4 distinct families depending on their biochemical and biophysical properties (Kinoshita 2003). Sept10 belongs to the Sept6 group (containing Sept6, Sept8, Sept10, Sept14, as well as Sept11, the latter also present in the Adgb interactome), which - unlike the other septin members - are catalytically inactive and constitutively bound to GTP (Sirajuddin et al. 2009). In sperm, Sept1, 4, 6, 7, and 12 have been localized to the sperm annulus, where they polymerize to a filamentous structure, forming a barrier between the midpiece and the principal piece of the spermatozoon (Toure et al. 2011). In this study, we focused on Sept10, and detected a strong signal in acrosomal granules in S7 spermatids, in the migrating annulus of S12 spermatids as well as in the annulus and neck region of mature sperm together with Adgb, suggesting a very early interaction between the two proteins, resulting in a missing signal (likely the annulus) upon absence of Adgb in knockouts. It is noteworthy that two independently generated mouse models for Sept4^-/-^ as well as Sept12^+/-^ mice are male infertile and harbour disorganized sperm mitochondria (Ihara et al. 2005, Kissel et al. 2005, Lin et al. 2009). Moreover, Sept4^-/-^ mice display a bent sperm tail and absence of annulus (Ihara et al. 2005, Kissel et al. 2005), whereas Sept12^+/-^ mice exhibit broken acrosomes, misshaped nuclei and increased apoptosis of germ cells (Lin et al. 2009). Correspondingly, SEPT12 mutations have been described in infertile men displaying abnormal sperm including defective annulus with a bent tail (Kuo et al. 2012). Additionally, defective sperm head morphology and DNA integrity have recently been reported for two different SEPT14 missense mutations (Wang et al. 2019, Lin et al. 2020). Previous findings proposed a septin ring assembly as octameric filaments at the annulus, classically composed of septins 12-7-6-2-2-6-7-12 or 12-7-6-4-4-6-7-12, in which septins 12 and 7 and septins 6 and 2/4 are connected through their GTP-binding domain (G-interface), whereas septins 7 and 6 and the two central septins 2/4 interact via their NC-termini (NC-interface) (Sirajuddin et al. 2007, Kuo et al. 2015). It was further proposed that septins from the same sub-group can substitute each other (Sellin et al. 2011). It is therefore tempting to speculate that Sept10 might replace Sept6 in the ring, thus Adgb’s localisation at the annulus. The identification of Sept7 and Sept2 as additional Adgb-interacting proteins is in favour of this assumption. Interestingly, another ring-like septin structure was recently described at the sperm neck, composed of Sept12 which complexes together with Sept1, Sept2, Sept10 and Sept11. Two mutations of Sept12 identified in patients disrupted the complex, leading to unstable head-tail junctions and defective connecting piece formation. Strikingly, the mutation of Sept12 and the subsequent disruption of the complex led to loss of Sept10 signal in the annulus (Shen et al. 2020) as also observed in Adgb-deficient mice. Accordingly, our data suggest an interdependence between Adgb and Sept10, which is required for the maintenance of the annulus, head shaping and proper mitochondrial localisation.

Septins have also been identified as components of cilia, which are hair-like microtubular protrusions at the surface of various cell types. Sept2 forms a diffusion barrier at the base of the cilium, impeding ciliary formation through loss of Sonic Hedgehog signalling when depleted (Hu et al. 2010). Sept2/7/9 form a complex that associates with the ciliary axoneme, thereby regulating ciliary length (Ghossoub et al. 2013). Accordingly, septin association with the cytoskeleton and particularly with microtubular structures has been extensively studied (Spiliotis and Nakos 2021), and numerous other cilia-related proteins participate in sperm flagellum formation.

Furthermore, most ciliopathies include male infertility and immotile sperm due to defective axonemal organisation (Brown and Witman 2014). Likewise, Adgb knockout mice display aberrant microtubule arrangements, thus it cannot be excluded that a scaffolding and simultaneous regulatory action between Adgb and Sept10 is necessary to support microtubular structure. Moreover, a recent study reported the consistent presence of Adgb in the ciliomes of three distinct evolutionary ancestral taxa, further suggesting a conserved function related to microtubular organisation and likely flagella formation (Sigg et al. 2017). In line with this study, our recent *in vitro* investigations have demonstrated that Adgb is transcriptionally regulated by FoxJ1, a master regulator of ciliogenesis (Koay et al. 2021).

A robust Adgb-Sept10 interaction was also detected by co-immunoprecipitation experiments, both in testis lysates as well as in HEK293 cells overexpressing ADGB and SEPT10. Unexpectedly, interaction was still observed upon truncating the N-terminal and C-terminal portions of ADGB. Furthermore, specific mutation of the coiled-coil domain of either ADGB or SEPT10 did not disrupt the interaction, suggesting that interaction of ADGB and SEPT10 occurs at various locations along the two proteins and that the proteins may entangle around each other. This close interaction may serve as targeting mechanism for the proteolytic processing of SEPT10 that we could observe upon ectopic expression of both proteins. The unique chimeric domain structure of Adgb with an N-terminal calpain-like protease domain and an IQ motif within a rearranged globin domain suggests a CaM-mediated regulation. Indeed, we could demonstrate that ADGB interacts with CaM via its IQ binding motif, and that the IQ binding motif seems pivotal in the observed proteolytic cleavage of SEPT10. CaM is a versatile protein, which can interact with a broad range of target proteins and act in a wide variety of cellular pathways and processes (Villalobo et al. 2018). Calmodulin-bound calcium represents a crucial activator of proteolytic activity of Ca^2+^-dependent calpain proteases (Villalobo, González-Muñoz, and Berchtold 2019, Bähler and Rhoads 2002). Most interestingly, calpain-dependent proteolytic cleavage of septins is not unprecedented: it was shown that Sept5 is a substrate of both calpain-1 and calpain-2 in platelets, where the cleavage triggers secretion of chemokine-containing granules (Randriamboavonjy et al. 2012). Additionally, it was reported that small modifications in septin structures may have profound consequences on the protein’s function. For example, a single point mutation within the N-terminus of Sept2 leads to homomeric filament formation without including any other septin type (Kim et al. 2012). Furthermore, the PKA-dependent phosphorylation of Sept12 at a single site leads to the disruption of the Sept12 filament, which dissociates from Sept7-6-2 and Sept7-6-4 complexes (Shen et al. 2017). Finally, SUMOylation failures are associated with aberrant filament formation for Sept6, 7 and 11, which are still able to interact with other endogenous septin members, remarkably in a stronger way than when properly SUMOylated (Ribet et al. 2017). It is therefore conceivable that CaM-dependent Adgb-mediated proteolytic cleavage of Sept10 may be a prerequisite for proper Sept10 function or localization within the sperm neck or annulus.

The presence of an oxygen-binding globin might be beneficial in a strongly hypoxic environment such as the testis. Indeed, testicular interstitial oxygen levels are very low, and tissue oxygen pO_2_ is estimated to represent only 20% of the testicular artery pO_2_ (Free, Schluntz, and Jaffe 1976), whereby the microvasculature controls the oxygen supply to the testicular tissue and thereby indirectly also controlling the oxygen reaching the large seminiferous tubules by diffusion, only (Reyes et al. 2012). We previously demonstrated heme hexa-coordination of Adgb (Bracke, Hoogewijs, and Dewilde 2018b), a unique characteristic also reported for mammalian Cygb and Ngb, rather associated with functionality beyond canonical oxygen transport and storage. Similar to the postulated cytoprotective functions against ROS of Cygb and Ngb (Burmester and Hankeln 2009, Burmester and Hankeln 2014, Randi et al. 2020, Keppner et al. 2020), it is tempting to speculate that Adgb might be involved in detoxification of harmful ROS as male infertility is affected by ROS, and spermatozoa are particularly sensitive to ROS-induced damage. The lack of Adgb-dependent differential regulation of redox sensitive genes on transcriptome or single gene level between wildtype and Adgb-deficient bulk testis tissue lysates argues against, but does not exclude, an anti-oxidative function of Adgb. In line with these observations, the absence of any O_2_-dependent effect of our *in vitro* findings on ADGB-dependent SEPT10 cleavage suggests that the proteolytic activity of Adgb is independent of a functional globin domain. Although we reported heme incorporation on recombinantly expressed human protein (Bracke, Hoogewijs, and Dewilde 2018b), Adgb orthologs in more basal organisms lack the crucial proximal His, suggesting that heme-binding might not be the most prominent characteristic feature of Adgb. Establishing a mechanistic explanation for the chimeric domain structure of the Adgb protein will remain a major challenge for the future.

In conclusion our study is the first to demonstrate a functional role for androglobin, the fifth mammalian globin. We present convincing *in vivo* evidence that Adgb is required for murine spermatogenesis. Interdependence between Adgb and Sept10 is necessary for sperm head shaping via the manchette and proper flagellum formation. *In vitro* data demonstrated CaM binding to ADGB and suggested that ADGB contributes to proteolytical cleavage of SEPT10 in a CaM-dependent manner. Our work provides a crucial contribution to the characterization of the physiological role of this novel enigmatic chimeric globin type.

## Materials and methods

### Animals, ethics statement and genotyping

All experimental procedures and animal maintenance followed Swiss federal guidelines and the study was revised and approved by the “*Service de la sécurité alimentaire et des affaires vétérinaires*” (SAAV) of the canton of Fribourg, Switzerland (license number 2017_16_FR). Animals were housed in rooms with a 12 hour/12 hour light/dark cycle, controlled temperature and humidity levels, and had free access to food and water.

Interbreeding of heterozygous animals was performed to obtain wildtype (+/+), heterozygous (Tg/+ for tm1a and tm1b, or +/- for tm1d), and homozygous/knockout (Tg/Tg for tm1a and tm1b, or -/- for tm1d) littermates, that were experimentally used, if not otherwise stated, between 3 to 9 months of age. Genotyping of tm1a and tm1b animals was performed using the following primers (**Fig. S1**): F1 5’**-** CCGTGCCCAGCTATATGAGT-3’; R1 5’-CACAACGGGTTCTTCTGTTAGTCC-3’; R2 5’-CCAGCGGTGTTCCTTTCTTA-3’. Primers for tm1d genotyping were the following (**Fig. S1**): F1, R2, and R4 5’**-**ACTGATGGCGAGCTCAGACC-3’. PCR amplification was performed for 36 cycles of 1 min at 95°C, 1 min at 56°C and 1 min at 72°C. The PCR products were separated by electrophoresis on 2% agarose gels and visualized by ethidium bromide staining.

### Gene targeting and knockout mouse generation

The Adgb^tm1a(KOMP)Wtsi^ (tm1a) strain was generated by blastocyst microinjection of embryonic stem (ES) cell clone EPD0707_3_H06, provided by the Knockout Mouse Project (KOMP) (Skarnes et al. 2011). Correct targeting of the Adgb locus was verified prior to microinjection by long-range PCR using primers 5’S 5’-CTGTACACTGGTTGTACACTGGTACAACTG-3’; 5’AS 5’-GGACTAACAGAAGAACCCGTTGTG-3’; 3’S 5’-CACACCTCCCCCTGAACCTGAAAC-3’; 3’AS 5’-GTACTTGATTGGACGATGATCCAAG-3’ (**Fig. S1**), generating a band of 6.7 kb for 5’ primers, and 5.1 kb for 3’ primers (**Fig. S1**). Targeted clones were confirmed by Southern blot analysis using a hybridization probe that targets exon 13 (**Fig. S1**) revealing a band of 4.1 kb (wildtype) or 3.2 kb (tm1a allele) following digestion of genomic DNA with *PvuII*, and a band of 2.8 kb (wildtype) or 2.4 kb (tm1a allele) following digestion of genomic DNA with *PstI* (**Fig. S1**). Chimeric mice were bred with C57BL/6-Tyr^c-Brd^ mice, and germline transmission in the F1 offspring was verified by PCR using primers F1, R1 and R2 (**Fig. S1**). The mice were further bred to C57BL/6N-Hprt^Tg(CMV-cre)Brd/Wtsi^ transgenic mice expressing the Cre allele to delete exons 13 and 14 and the neo cassette to generate the Adgb^tm1b(KOMP)Wtsi^ strain (tm1b), or to C57BL/6N-Gt(ROSA)26Sor^tm1(FLP1)Dym/Wtsi^ transgenic mice expressing the Flp recombinase to delete the whole transgene cassette, thereby generating the Adgb^tm1c(KOMP)Wtsi^ strain. The latter were further crossed with C57BL/6N-Hprt^Tg(CMV-cre)Brd/Wtsi^ transgenic mice to delete exons 13 and 14, thereby generating the Adgb^tm1d(KOMP)Wtsi^ (tm1d, knockout) strain. The Cre recombinase allele was bred out before any experiments were performed.

### RNA extraction and RT-qPCR

Testes were frozen in liquid nitrogen and stored at −80°C. Tissues were homogenized using a TissueLyser (Qiagen, Valencia, CA, USA). Subsequent RNA isolation and cDNA synthesis were performed as described previously (Keppner et al. 2019). In brief, RNA was extracted using an RNeasy Mini Kit (Qiagen,) and reverse-transcription (RT) was performed with 1.5 µg of total RNA and PrimeScript reverse transcriptase (Takara Bio Inc, Kusatsu, Japan). RT-quantitative (q)-PCR was performed on a CFX96 C1000 real-time PCR cycler (Bio-Rad Laboratories, Hercules, CA) using SYBRgreen PCR master mix (Kapa Biosystems, London, UK). 21.5 ng of cDNA were loaded, and each sample was run as duplicate. mRNA levels were normalized to β-actin as previously described (De Backer et al. 2021). Primer sequences are displayed in **Table S1**.

### Cell culture and transfection

HEK293 and A375 (ATCC CRL-1619) cells were maintained in Dulbecco’s Minimum Essential Media (DMEM) (Gibco, Life Technologies, Carlsbad, CA, USA), containing L-glutamine, supplemented with 10% heat-inactivated fetal bovine serum FBS (PAN Biotech, Aidenbach, Germany) and 100 Units/mL penicillin/100 μg/mL streptomycin (Gibco, Life Technologies, Carlsbad, CA, USA). Both cell lines were incubated in a humidified 5% CO_2_ atmosphere at 37°C and were routinely subcultured after trypsinization. For hypoxic experiments, cells were seeded out in 6-well plates or 100-mm culture dishes. The subsequent day, hypoxia experiments were carried out at 0.2% O_2_ and 5% CO_2_ in a gas-controlled glove box (InvivO2 400, Baker Ruskinn, Bridgend, UK) for 24 hours. Transfection of HEK293 cells was performed using calcium-phosphate (Jordan, Schallhorn, and Wurm 1996), with 2 µg of plasmid DNA for regular immunoblotting experiments, and 5 µg of plasmid DNA for immunoprecipitation experiments. Briefly, the DNA was diluted in sterile water and mixed with 250 mM CaCl_2_. 25 µM chloroquine was added to the cells and allowed to incubate for a minimum of 20 min. Prewarmed 37°C 2× HBS buffer pH 7.05 (NaCl 280 mM, KCl 10 mM, Na_2_HPO_4_ 1.5 mM, D-glucose 12mM, HEPES 50mM) was added to the DNA solution (50% v/v), and the transfection mixture was added dropwise to the cells. The medium was replaced after 6 hours. Transfection of A375 cells was performed using JetOptimus (Polyplus-transfection SA, Illkirch-Grafffenstaden, France) according to the manufacturer’s instructions.

### SDS-PAGE and immunoblotting

Tissues were homogenized as described (Keppner et al. 2019). Cells were lysed in triton buffer (Tris-HCl [20 mM, pH 7.4], NaCl [150 mM], triton X-100 [1%]), left on ice for 15 min, centrifuged, and the proteins were quantified by Bradford assay. 25 µg of proteins were separated by SDS-PAGE on 10% gels, and proteins were electrotransferred to nitrocellulose membranes (Amersham Hybond-ECL, GE Healthcare, Chicago, IL, USA). The membranes were incubated overnight at 4°C with primary antibody (**Table S2**) and for 1 hour with donkey anti-rabbit or anti-mouse IgG HRP-conjugated secondary antibody (1:5000, Amersham, Bukinghampshire, UK). All antibodies were diluted in TBS-tween (1%) and dried milk (1%). The signal was revealed using ECL Prime (Amersham, Bukinghampshire, UK) on a C-DiGit Western blot scanner (LI-COR Biosciences), and quantified using ImageStudio program (LI-COR Biosciences, Lincoln, NE, USA). The polyclonal anti-Adgb antibody was custom-made (Proteintech Group Inc, Rosemont, IL, USA). A fusion protein immunogen raised against the 409-745 amino acid region of mouse Adgb was used for the immunisation of two rabbits over a period of 102 days. The antibodies in immune sera were affinity-purified. Pre-bleeds, test bleeds and purified antibodies were tested and validated by immunoblotting on wildtype and knockout testis extracts.

### Immunoprecipitation

For immunoprecipitation (IP) of subsequent LC-MS/MS and immunoblotting analyses, 4 and 2 mg of proteins were used, respectively. The protein lysates were first pre-cleared for 24 hours at 4°C with G-sepharose beads (GE Healthcare, Chicago, IL, USA) coupled to rabbit IgG (Bethyl Laboratories Inc, Montgomery, TX, USA). Samples were incubated overnight at 4°C with 2 µg primary antibody (**Table S2**) or 2 µg rabbit IgG, followed by 4 hours with G-sepharose beads, then washed 2 times with wash buffer (Tris-HCl [20 mM, pH 7.4], NaCl [300 mM], LAP [1 mM]), and 3 times with equilibration buffer (Tris-HCl [20 mM, pH 7.4], NaCl [150 mM], LAP [1 mM]). Samples were eluted by boiling for 5 min at 95°C in 2× sample buffer, and separated from the beads by centrifugation.

### LC-MS/MS analysis

Washed IP beads were incubated with *Laemmli* sample buffer and proteins were reduced with 1 mM DTT for 10 min at 75°C and alkylated using 5.5 mM iodoacetamide for 10 min at room temperature. Protein samples were separated by SDS-PAGE on 4-12% gradient gels (ExpressPlus, Genscript, New Jersey, NJ, USA). Each gel lane was cut into 6 equal slices, the proteins were in-gel digested with trypsin (Promega, Madison, WI, USA), and the resulting peptide mixtures were processed on StageTips (Rappsilber, Mann, and Ishihama 2007, Shevchenko et al. 2006).

LC-MS/MS measurements were performed on a Q Exactive Plus mass spectrometer (Thermo Fisher Scientific, Waltham, MA, USA) coupled to an EASY-nLC 1000 nanoflow HPLC (Thermo Fisher Scientific). HPLC-column tips (fused silica) with 75 µm inner diameter were packed with Reprosil-Pur 120 C18-AQ, 1.9 µm (Dr. Maisch GmbH, Ammerbuch, Germany) to a length of 20 cm. A gradient of solvents A (0.1% formic acid in water) and B (0.1% formic acid in 80% acetonitrile in water) with increasing organic proportion was used for peptide separation (loading of sample with 0% B; separation ramp: from 5-30% B within 85 min). The flow rate was 250 nL/min and for sample application 650 nL/min. The mass spectrometer was operated in the data-dependent mode and switched automatically between MS (max. of 1×10^6^ ions) and MS/MS. Each MS scan was followed by a maximum of ten MS/MS scans using normalized collision energy of 25% and a target value of 1000. Parent ions with a charge state form z = 1 and unassigned charge states were excluded from fragmentation. The mass range for MS was m/z = 370-1750. The resolution for MS was set to 70,000 and for MS/MS to 17,500. MS parameters were as follows: spray voltage 2.3 kV; no sheath and auxiliary gas flow; ion-transfer tube temperature 250°C. The MS raw data files were uploaded into the MaxQuant software version 1.6.2.10 for peak detection, generation of peak lists of mass error corrected peptides, and for database searches (Tyanova, Temu, and Cox 2016). A full-length UniProt mouse (based on UniProt FASTA version April 2016) or human database (UniProt FASTA version March 2016) additionally containing common contaminants, such as keratins and enzymes used for in-gel digestion, was used as reference. Carbamidomethylcysteine was set as fixed modification and protein amino-terminal acetylation and oxidation of methionine were set as variable modifications.

Three missed cleavages were allowed, enzyme specificity was trypsin/P, and the MS/MS tolerance was set to 20 ppm. The average mass precision of identified peptides was in general less than 1 ppm after recalibration. Peptide lists were further used by MaxQuant to identify and relatively quantify proteins using the following parameters: peptide and protein false discovery rates, based on a forward-reverse database, were set to 0.01, minimum peptide length was set to 7, minimum number of peptides for identification and quantitation of proteins was set to one which must be unique. The ‘match-between-run’ option (0.7 min) was used.

### Propidium iodide staining and flow cytometry

Preparation of germ cell suspensions was achieved as described (Jeyaraj, Grossman, and Petrusz 2003). Briefly, decapsulated testes were incubated in 0.5 mg/mL collagenase type IV in PBS, washed with PBS, and incubated in 1 µg/mL DNase and 1 µg/mL trypsin. Soybean trypsin inhibitor was added, the suspension was filtered, washed in PBS, fixed with 70% ethanol and stored at 4°C. DNA staining using propidium iodide was performed as described (Krishnamurthy et al. 2000). Propidium iodide-stained cells were analysed in a FACScan flow cytometer (Becton-Dickinson Immunocytometry, San Jose, CA, USA). Cell populations were selected based on their DNA content, and their relative numbers were calculated using Summit (Cytomation, CO, USA).

### Histological analyses and immunofluorescence

Testes were fixed in 4% paraformaldehyde and embedded in paraffin. Preparation of sections and H&E staining was performed as described (Keppner et al. 2015). Pictures were taken using a Nikon Eclipse microscope (Nikon Corporation, Tokyo, Japan).

For immunofluorescence, testes were fixed in 4% paraformaldehyde (PFA) for at least 1 week, and incubated in 30% sucrose for another week. The testes were embedded in Optimal Cutting Temperature compound (O.C.T. Tissue-Tek, Sakura Finetek, Tokyo, Japan), and 5 µm thick sections were cut on a cryotome. For seminiferous tubule dissections and stainings, slides were prepared as previously described (Kotaja et al. 2004). For sperm stainings, cauda epididymal sperm was retrieved and diluted in PBS. A drop of the suspension was smeared on glass sections, and fixed by drying for 15 min and by 4% PFA for 20 min. The slides were blocked in 10% normal goat serum and 0.5% triton X-100 for 1 hour. Testis sections and sperm slides were incubated overnight at 4°C with primary antibodies (**Table S5**) in 5% normal goat serum and 0.25% triton X-100. The slides were washed with PBS (3×10 min), and incubated with secondary Alexa Fluor 488 or 594 coupled goat anti-mouse or anti-rabbit IgG (1:300, Invitrogen, Waltham, MA, USA) for 1 hour, washed again with PBS (3×10 min), and counterstained with Sudan Black (0.1%) for autofluorescence quenching. Slides were mounted with fluoromount mounting medium containing DAPI (SouthernBiotech, Birmingham, AL, USA), and visualized using a Nikon Eclipse fluorescent microscope (Nikon Corporation).

### In situ hybridization via RNAscope

RNAscope *in situ* hybridization was performed using BaseScope Detection Reagent Kit v2 RED (Advanced Cell Diagnostics Inc, Newark, CA, USA, Cat. No. 323900) according to the manufacturer’s instructions. H_2_O_2_ treatment, antigen retrieval and protease treatment were performed on 5 µm-thick sections prior to hybridization with probes for Adgb (BA-Mm-Adgb-3zz-st, Advanced Cell Diagnostics, Cat. No. 862141), DapB as negative control (BA-DapB-3zz, Advances Cell Diagnostics, Cat. No. 701011) and Ppib as positive control (Ba-Mm-Ppib-3zz, Advanced Cell Diagnostics, Cat. No. 701071) at 40°C for 2 hours followed by eight amplification steps. The signal was revealed with Fast Red, the sections were counterstained with Gill’s hematoxylin no. 1 and mounted with VectaMount (Vector Laboratories, Burlingame, CA, USA). The sections were visualized on a light microscope.

### Electron microscopy

All electron microscopy experiments were performed at the Electron Microscopy Platform of the University of Lausanne, Switzerland.

Mouse testes were fixed in 2.5% glutaraldehyde solution (EMS, Hatfield, PA, USA) in phosphate buffer (PB 0.1 M [pH 7.4]) for 1 hour at room temperature and postfixed in a fresh mixture of osmium tetroxide 1% (EMS) with 1.5% of potassium ferrocyanide (Sigma, St. Louis, MO, USA) in PB buffer for 1 hour at room temperature. The samples were then washed twice in distilled water and dehydrated in acetone solution (Sigma) at graded concentrations (30% - 40 min; 50% - 40 min; 70% - 40 min; 100% - 2×1 hour). This was followed by infiltration in in Epon resin (EMS, Hatfield, PA, USA) at graded concentrations (Epon 33% in acetone-4 hours; Epon 66% in acetone-4 hours; Epon 100%-2×8 hours) and finally polymerized for 48 hours at 60°C in an oven. Ultrathin sections of 50 nm thick were cut using a Leica Ultracut (Leica Mikrosysteme GmbH, Vienna, Austria), picked up on a copper slot grid 2×1 mm (EMS, Hatfield, PA, USA) coated with a polystyrene film (Sigma, St Louis, MO, USA). Sections were post-stained with uranyl acetate (Sigma, St Louis, MO, USA) 4% in H_2_O for 10 min, rinsed several times with H_2_O followed by Reynolds lead citrate in H_2_O (Sigma, St Louis, MO, USA) for 10 min and rinsed several times with H_2_O. Micrographs were taken with a transmission electron microscope FEI CM100 (FEI, Eindhoven, The Netherlands) at an acceleration voltage of 80kV with a TVIPS TemCamF416 digital camera (TVIPS GmbH, Gauting, Germany).

### RNA-seq library preparation and transcriptome sequencing

Total RNA from 2 independent samples of wildtype and Adgb^-/-^ testis was extracted using the mirVana miRNA-Kit according to manufacturer’s instructions (Life Technologies, Carlsbad, US). Prior to library construction, RNA quality was assessed using an Agilent 2100 Bioanalyzer and the Agilent RNA 6000 Nano Kit (Agilent Technologies, Santa Clara, CA, USA). RNA was quantified using Qubit RNA BR Assay Kit (Invitrogen, Waltham, MA, USA). Libraries were prepared starting from 1000 ng of total RNA using the RNA Sample Prep Kit v2 (Illumina Inc, San Diego, CA, USA) including a poly-A selection step following the manufacturer’s instructions and sequenced as 2 x 100 nt paired-end reads using an Illumina HiSeq 2500. Library preparation and sequencing were performed by the NGS Core Facility of the Department of Biology, Johannes-Gutenberg University (Mainz, Germany). RNA-Seq data are available at the European Nucleotide Archive under accession number PRJEB46499.

*Differential gene expression, GO term annotation and pathway enrichment analyses* Raw sequences were pre-processed to remove low quality reads and residual Illumina adapter sequences using BBduk from the BBtools suite (https://sourceforge.net/projects/bbmap/). The overall sequencing quality and the absence of adapter contamination were evaluated with FastQC. Mapping was performed with HISAT2 and quantification of gene expression was done using StringTie.

Differentially expressed genes were determined using DESeq2. Genes were considered differentially expressed when presenting |fold change| > 2 and false discovery rate (FDR)-corrected p-value ≤ 0.1. GO term enrichment analyses were performed using WebGestalt 2019 using the Overrepresentation Enrichment Analysis method, requiring a BH-corrected p-value ≤ 0.05 and a minimum enrichment of 4 genes for term/pathway.

Enrichment in Canonical Pathways were performed with Qiagen’s Ingenuity Pathway Analysis (IPA, Qiagen, Hilden, Germany), Core analysis tool using bias-corrected z-score (when applicable) and BH-corrected p-values ≤ 0.05.

### Cloning and construction of expression plasmids

Generation of pLenti6-ADGB was described before (Bracke, Hoogewijs, and Dewilde 2018b), pLenti6-SEPT10-V5 was purchased from DNASU (clone ID HsCD00943271, DNASU Plasmid Repository, Arizona State University, AZ, USA). All additional recombinant genes were cloned into pFLAG-CMV^TM^-6a expression vector (Sigma) unless otherwise specified. All genes were amplified by PCR using Phusion High-Fidelity DNA polymerase (Thermo Fisher Scientific). Recombinant *SEPT10* with N-terminal FLAG tag and C-terminal myc tag was constructed by amplifying and ligating SEPT10 coding sequence with in-primer designed myc tag (EQKLISEEDL) into the expression vector, in-frame with the N-terminal FLAG tag. A glycine-serine (GSG) linker was added between the last codon of *SEPT10* and the first codon of the myc tag. Truncated ADGB proteins consisting of the calpain-like domain, 350-residue uncharacterized domain and globin domain (N-terminal mutant), or the 700-residue uncharacterized region (C-terminal mutant) domain were designed with GFP tags at both N- and C-termini. The N-terminal mutant ADGB was amplified between codons of residues 58 – 968 in ADGB while the C-terminal mutant ADGB was amplified between codons of residues 969 – 1667. Amplicons were designed with 5’- and 3’-overhangs compatible with two customized GFP amplicons designed to anneal at the 5’- and 3’-ends of the genes. Glycine-serine linkers (GGSGGGGSGG) were added to bridge the GFP tags and the truncated ADGB proteins. Similarly, N-terminally GFP-tagged isolated ADGB globin was cloned by amplifying and ligating amplicons of the *ADGB* globin gene downstream to a *GFP* gene with complementary overhangs, with the glycine-serine linker added between the two proteins. With the same construction, GFP-tagged ADGB globin domains with a single mutation on the proximal histidine in helix F codon 8 (H824G) or the distal glutamine in helix E codon 7 (Q792G), or both (H824G/Q792G) were cloned by amplifying and ligating the globin gene using primers designed to carry the mutated codon sequence. ADGBΔIQ and ADGBΔCCD were constructed by amplifying designed *ADGB* amplicons with compatible overhangs and were ligated in-frame to generate a *ADGB* gene with deletions in the desired domains. For mammalian 2-hybrid assays, Gal4-CaM and VP16-globin domain were cloned into a pcDNA3.0 expression vector. Gal4 DNA-binding domain or VP16 transactivation domain sequences were amplified with the in-primer designed glycine-serine linker at the 3’-end of the amplicons. The coding sequence of *CALM3* and *ADGB* globin domain were amplified with complementary 5’-end and ligated to the Gal4 and VP16 sequences, respectively to generate the fusion genes.

### Reporter gene assays

For mammalian 2-hybrid assays 2.15 x 10^5^ HEK293 or 4 x 10^5^ A375 cells were transiently transfected with 1 µg firefly luciferase reporter plasmid (5xGAL4-TATA-luciferase, Addgene, 46756) (Sun et al. 1994) and 500 ng or 200 and 300 ng chimeric Gal4 and VP16 fusion protein vectors, respectively, in 12- or 6-well format using CaCl_2_ or JetOptimus. To control for differences in transfection efficiency and extract preparation, 25 ng or 50 ng pRL-SV40 *Renilla* luciferase reporter vector (Promega, Madison, WI, USA) was co-transfected, respectively for HEK293 and A375 cells.

Cultures were evenly split onto 12-well plates 24 hours after transfection for A375 cells. For hypoxia control experiments, 4 x 10^5^ A375 cells were transiently co-transfected with 500 ng firefly luciferase *EPO* reporter plasmid (Storti et al. 2014) and 50 ng pRL-SV40 *Renilla* luciferase reporter vector. Luciferase activities of duplicate wells were determined using the Dual Luciferase Reporter Assay System (Promega) as described before (Schörg et al. 2015). Reporter activities were expressed as relative firefly/*Renilla* luciferase activities. All reporter gene assays were performed at least 3 times independently.

### Testosterone quantification

Serum testosterone levels were determined as described previously with minor adaptations (Strajhar et al. 2016). Briefly, for solid-phase extraction (SPE), each serum sample (100 µL) was mixed with protein precipitation solution (100 µL, 0.8 M zinc sulphate in water/methanol; 50/50 v/v) containing 33 nM deuterium-labeled testosterone (D2) as internal standard. Prior SPE, all samples were diluted to a final volume of 1 mL with water and incubated in a thermoshaker (10 min at 4 °C, 1300 rpm). Following incubation, samples were centrifuged for 10 min at 16000 g at 4°C, and supernatants (950 µL) were transferred to Oasis HBL SPE (1 cc) cartridges (Waters, Milford, MA, USA), preconditioned with methanol and water (3 x1 mL each). Samples were washed with water (2 x 1 mL) and methanol/water (2 x 1 mL, 10/90 v/v). Testosterone was eluted with methanol (2 x 0.75 mL), evaporated to dryness (3 hours, 35°C) and reconstituted in methanol (25 µL, 10 min, 4°C, 1300 rpm). Testosterone content was analyzed by ultra-performance liquid chromatography-MS/MS (UPLC-MS/MS) using an Agilent 1290 Infinity II UPLC coupled to an Agilent 6495 triple quadrupole mass spectrometer equipped with a jet-stream electrospray ionization interface (Agilent Technologies). Analyte separation was achieved using a reverse-phase column (1.7 µm, 2.1 mm x 150 mm; Acquity UPLC BEH C18; Waters). Data acquisition and quantitative analysis was performed by MassHunter (Version B.10.0. Build 10.0.27, Agilent Technologies).

## Statistical analysis

All values were presented as mean ± standard error of the mean (SEM). Differences in means between two groups were analyzed with unpaired 2-tailed Student’s t-test (Fig. 1A,C,D,E; Fig. 6H right graph; Fig. S4B,C) and those among multiple groups with one-way ANOVA followed by Tukey posthoc test (Fig. 2A; Fig. 6D,G,H left graph). All statistics were performed with GraphPad Prism software 7.05. Values of p≤0.05 were considered statistically significant.

## Acknowledgements

We thank Christine Roulin for technical assistance. We thank Damien De Bellis from the Electron Microscopy Platform of the University of Lausanne for EM section preparation and image acquisition. This work was supported by the Swiss National Science Foundation to DH (grant 31003A_173000) and the German Research Foundation to DH (HO 5837/1-1) and TH (HA 2103/9-1).

## Supplementary Information

**Figure S1.**
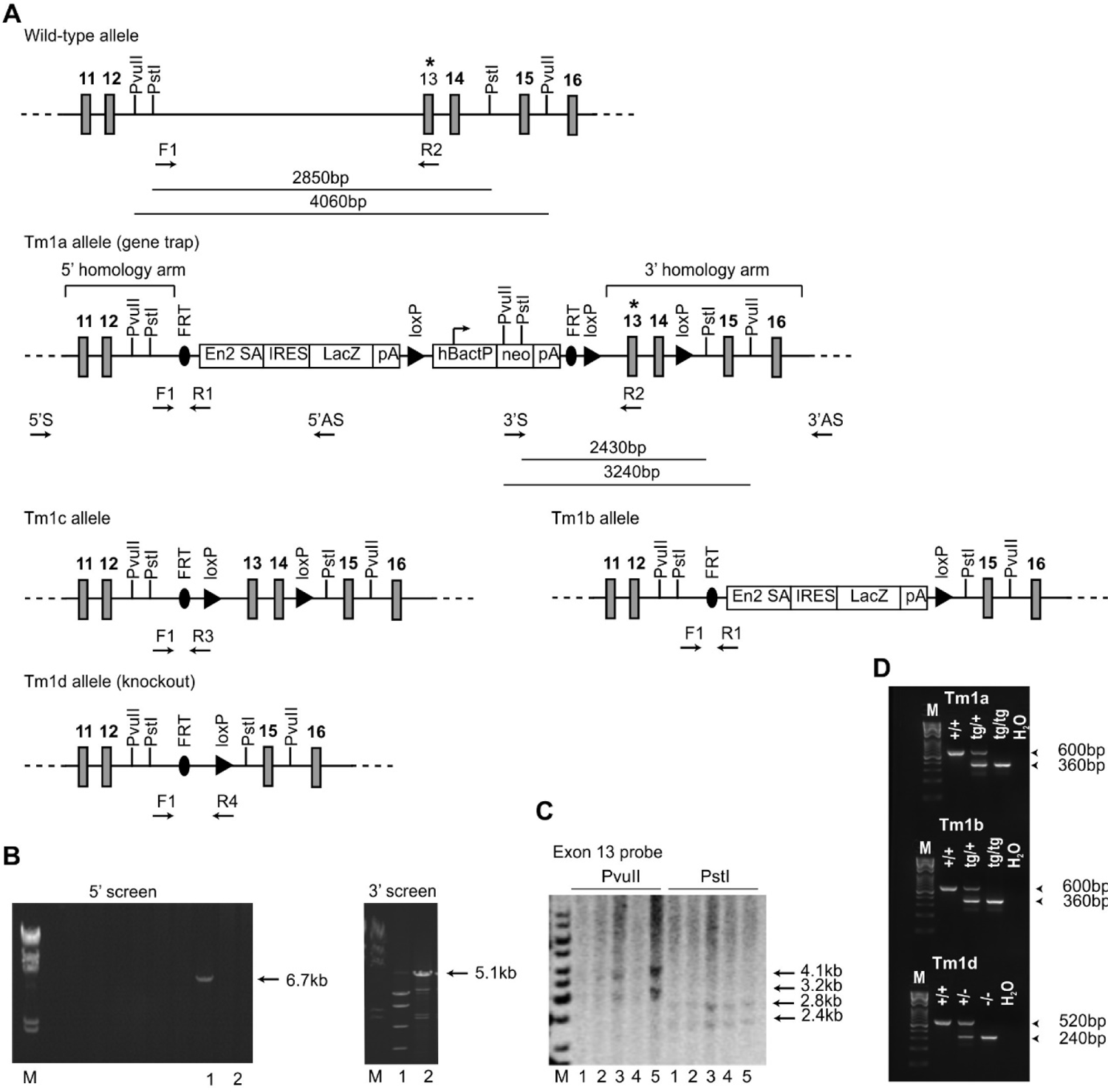
Generation of Adgb knockout mice. (**A**) Schematic representation of the wild-type, Tm1a (gene-trap), Tm1b (following breeding with Cre-deleter mice), Tm1c (following breeding with Flp-deleter mice) and Tm1d (knockout) (following breeding of Tm1c mice with Cre-deleter mice) alleles. The position of the probe (asterisk), the primers (5’S, 5’AS, 3’S and 3’AS) and restriction sites for the ES cell screening by PCR and Southern blot are shown. FRT sites flank the gene-trap construct, containing the lacZ gene and the neomycin resistance cassette (neo), whereas loxP sites flank the neo-cassette and exons 13 and 14 of the *Adgb* gene. Position of the PCR primers for the genotyping of the different mouse lines are shown (F1 and R1-4). (**B**) Representative PCR-based analysis of targeted ES cells using primers 5’S and 5’AS (left panel) (M: marker, 1 and 2: positive and negative clones respectively), and primers 3’S and 3’AS (right panel) (M: marker, 1 and 2: negative and positive clones respectively). (**C**) Southern blot analysis of targeted ES cell clones using the exon 13 probe (asterisk) and following digestion with *PvuII* and *PstI* (M: marker, 1-5: different clones tested positive for both 5’ and 3’ PCR reactions). (**D**) Genotyping of Tm1a wild-type (+/+), heterozygous (tg/+) and homozygous (tg/tg) mice (upper panel), M: marker, Tm1b wild-type (+/+), heterozygous (tg/+) and homozygous (tg/tg) mice (middle panel), and Tm1d wild-type (+/+), heterozygous (+/-) and homozygous (-/-, knockout) mice (lower panel).

**Figure S2.**
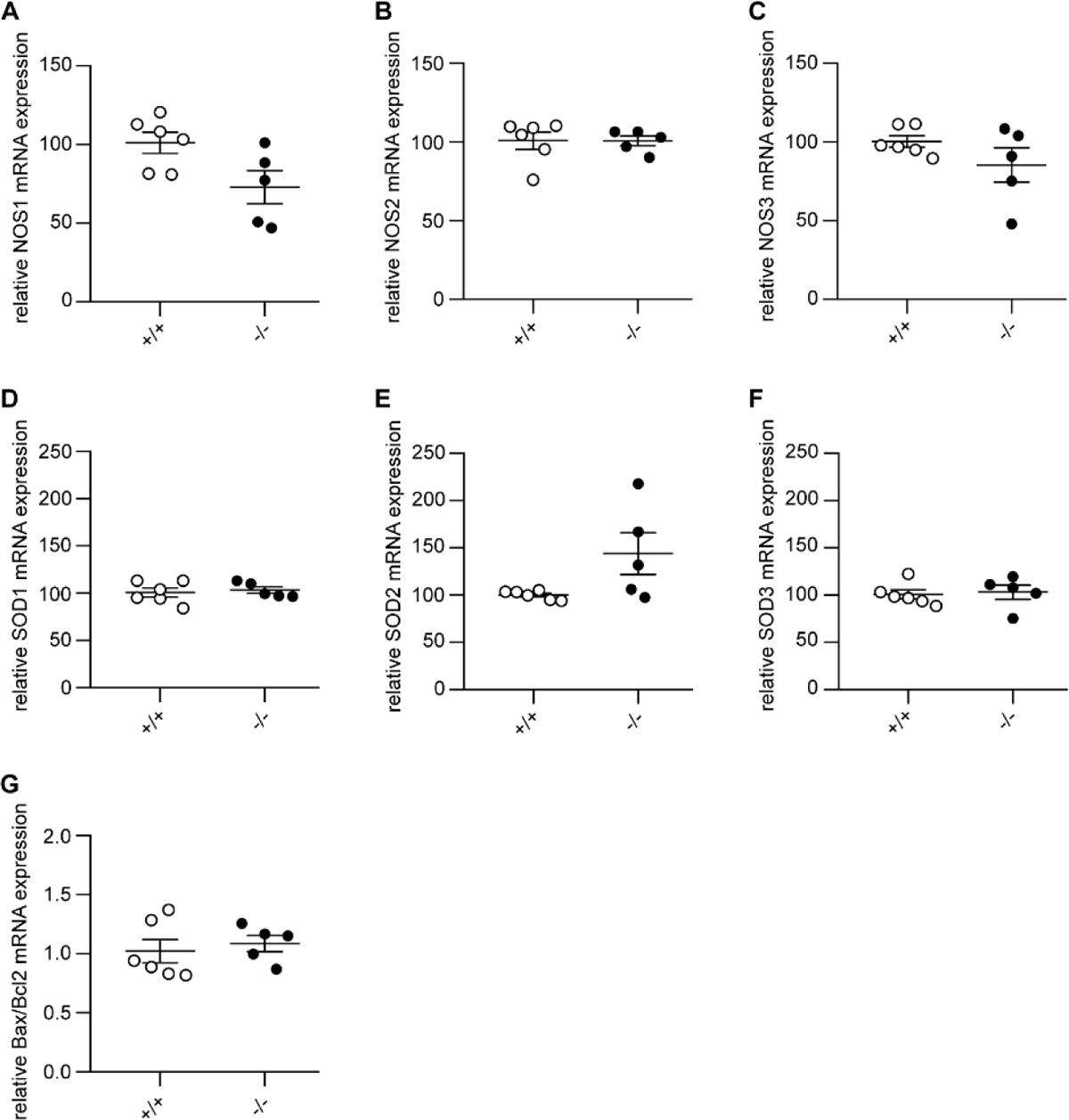
Adgb KO mice do not display changes in NOS, SOD and apoptotic gene expression. Relative mRNA expression levels of (**A**) NOS1, (**B**) NOS2, (**C**) NOS3, (**D**) SOD1, (**E**) SOD2, (**F**) SOD3, and (**G**) ratio of Bax and Bcl2 in testis lysates from Adgb wildtype (+/+, n=6) and knockout (-/-, n=5) mice.

**Figure S3.**
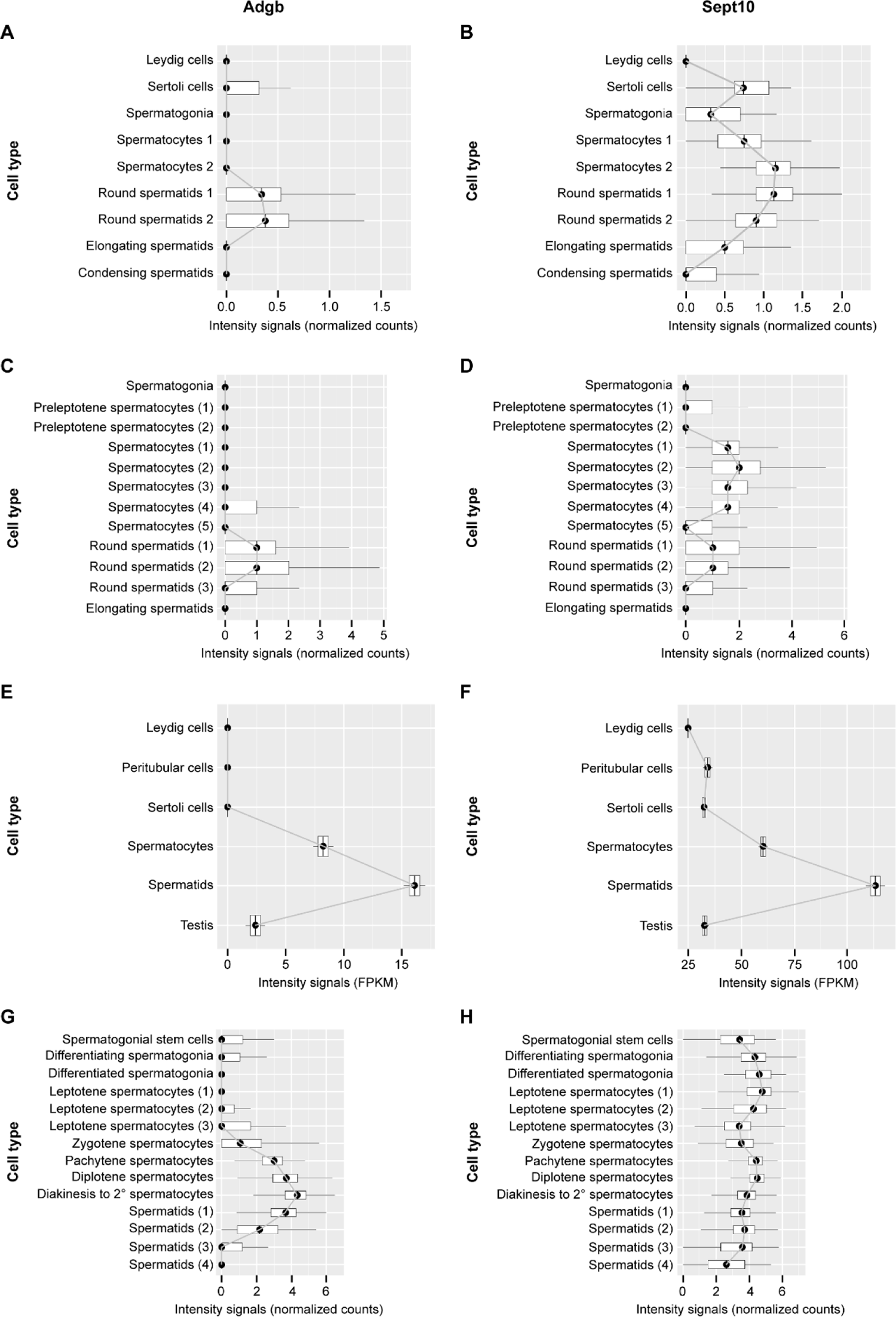
Temporal Adgb and Sept10 expression profiles based on single-cell and bulk RNA-seq datasets. **(A, C, E, G)**: Adgb; **(B, D, F, H)**: Sept10. Data were obtained from Lukassen et al. 2018 (mouse, panels A and B), Green et al. 2018 (mouse, panels C and D), Jégou et al. 2017 (human, panels E and F), and Wang et al. 2018 (human, panels G and H), respectively (Green et al. 2018, Jégou et al. 2017, Lukassen et al. 2018, Wang et al. 2018).

**Figure S4.**
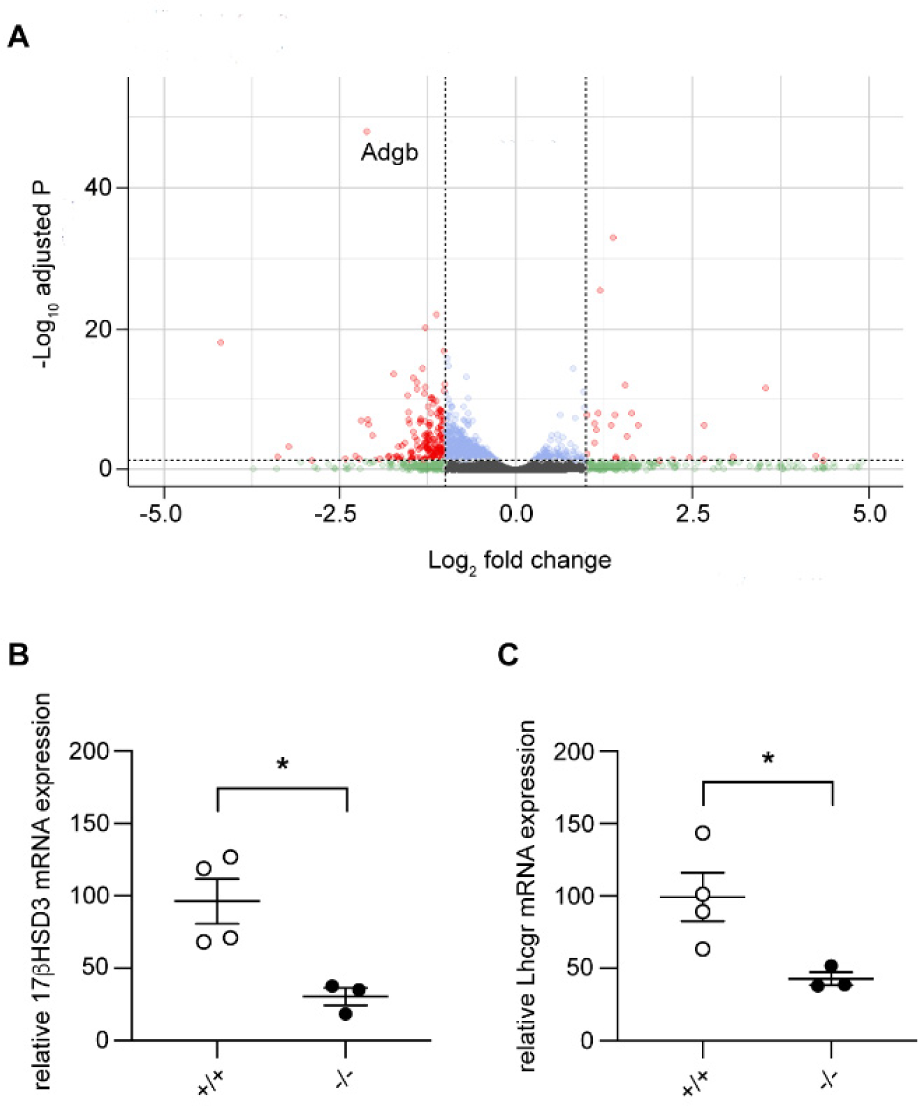
Volcano plot of differentially expressed genes in Adgb knockout mice testis samples and validation by RT-qPCR. (A) X-axis represents the log2 fold change and y-axis represents log10 of the adjusted P-value. Genes are assigned with specific colors after DESeq2 analysis: gray (not significant [NS]), green |Log2FC|>1, blue (adjusted P<0.05), or red (|Log2FC|>1 and adjusted P<0.05). Adgb is indicated. **(B)** Relative mRNA expression levels of 17βhds3 in testes of wildtype (+/+) and knockout mice (-/-) (n=3-4 per genotype) on post-natal day 24. (**C**) Relative mRNA expression levels of Lhcgr in testes of wildtype (+/+) and knockout mice (-/-) (n=3-4 per genotype) on post-natal day 24. * p< 0.05.

**Figure S5.**
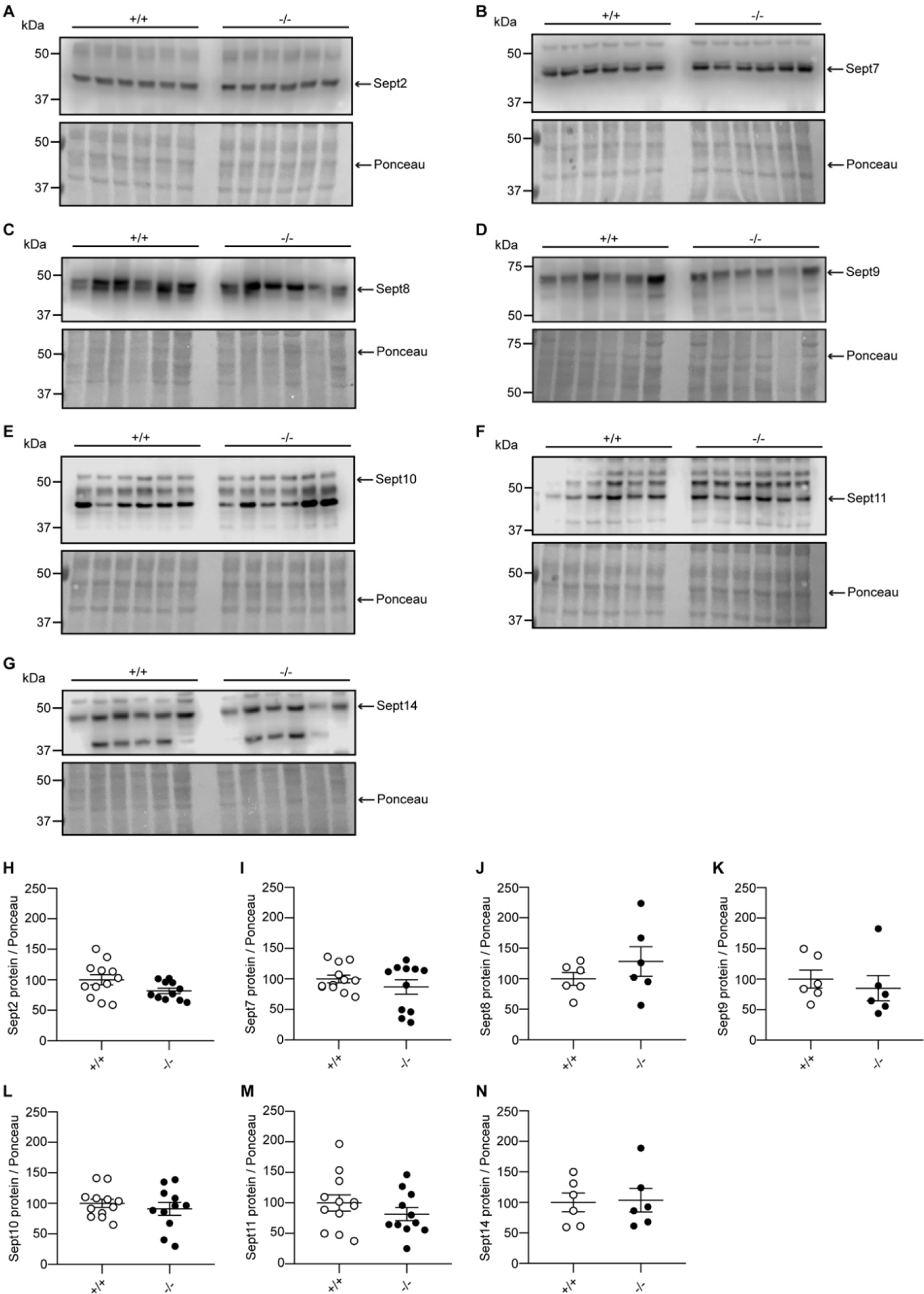
The protein expression levels of Sept2, 7, 8, 9, 10, 11 and 14 are unaffected in Adgb KO testis. Representative immunoblots of testis lysates from Adgb wildtype (+/+, n=6-12) and knockout (-/-, n=6-11) mice for (**A**) Sept2, (**B**) Sept7, **(C)** Sept8, (**D**) Sept9, (**E**) Sept10, (**F**) Sept11, and (**G**) Sept14, and the corresponding protein quantifications for (**H**) Sept2, (**I**) Sept7, (**J**) Sept8, (**K**) Sept9, (**L**) Sept10, (**M**) Sept11, and (**N**) Sept14. Ponceau S protein staining was used as loading control.

**Figure S6.**
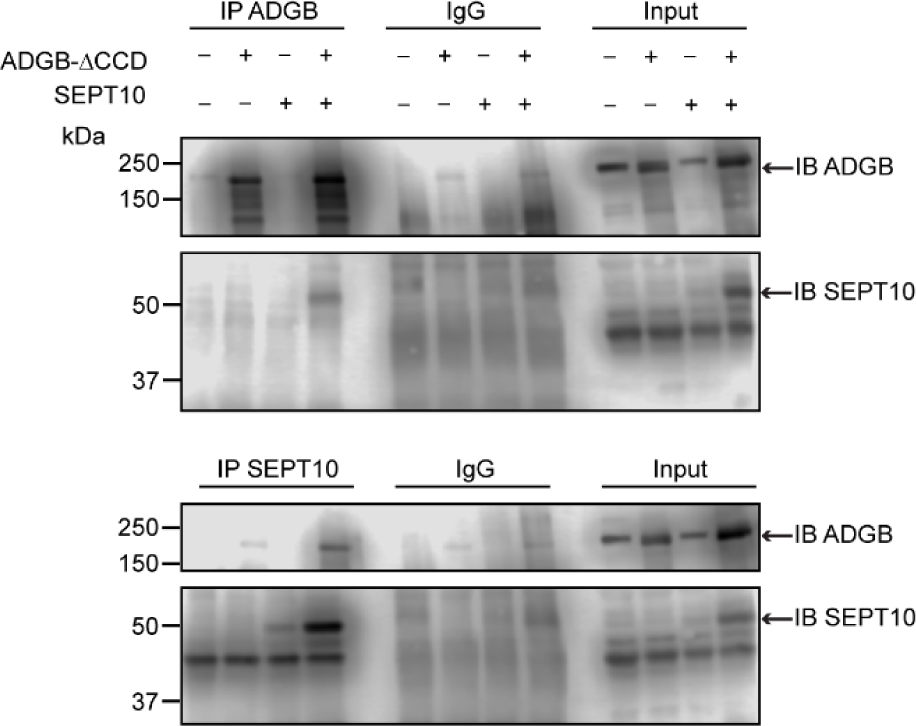
The interaction between ADGB and SEPT10 is maintained despite mutation of the coiled-coil domains. Representative immunoblots of ADGB and Sept10 in protein lysates of HEK293 cells (co-)transfected with CCD mutant ADGB and CCD mutant SEPT10 following co-immunoprecipitation of ADGB and SEPT10.

**Figure S7.**
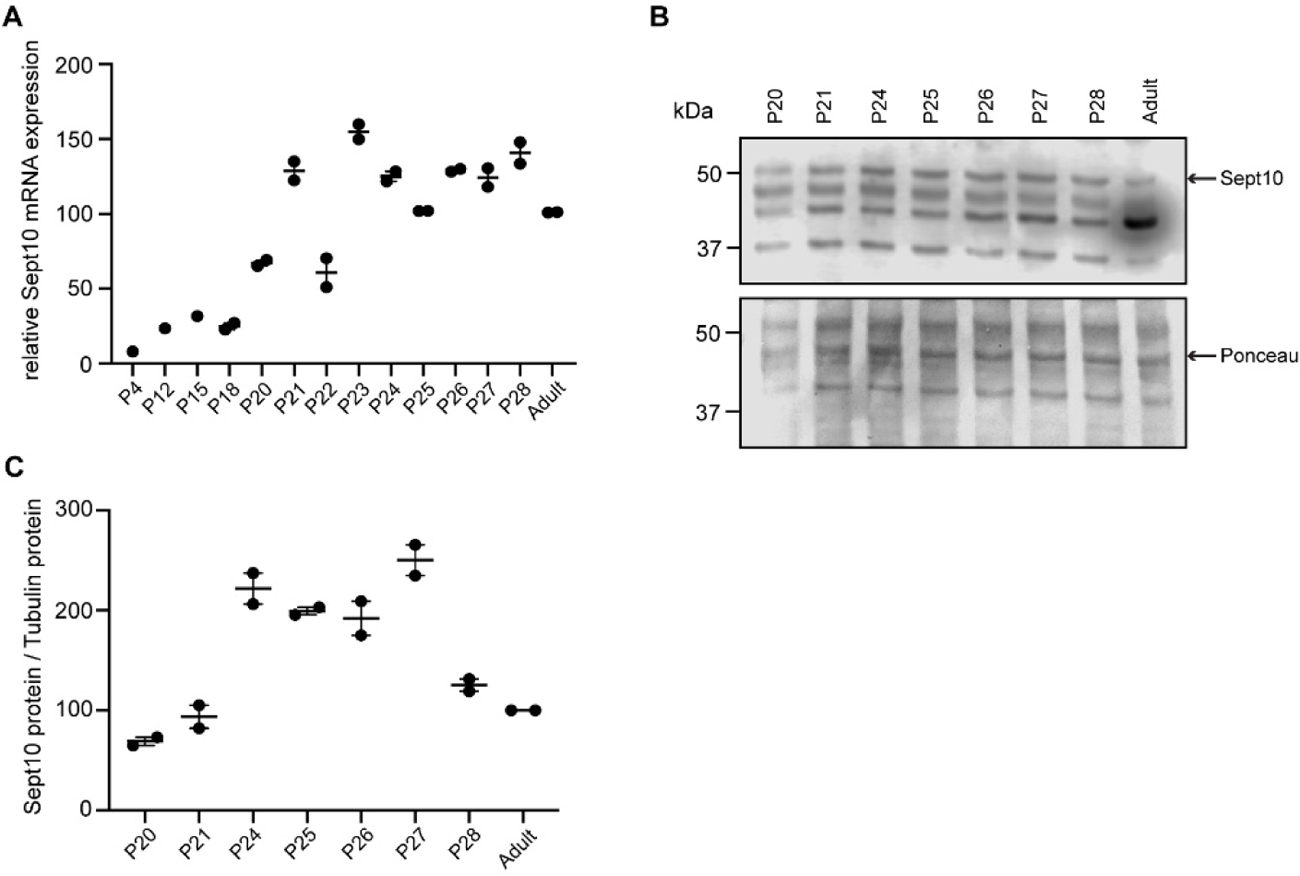
Sept10 temporal expression profile on mRNA and protein levels. (**A**) Relative mRNA expression levels of Sept10 in testes of wildtype mice during the first wave of spermatogenesis at indicated post-natal (P) days. (**B**) Representative immunoblot for Sept10 in testis lysates from wildtype mice at indicated post-natal (P) ages (n=2 per condition) and (**C**) corresponding protein quantification. Ponceau S protein staining was used as loading control.

**Figure S8.**
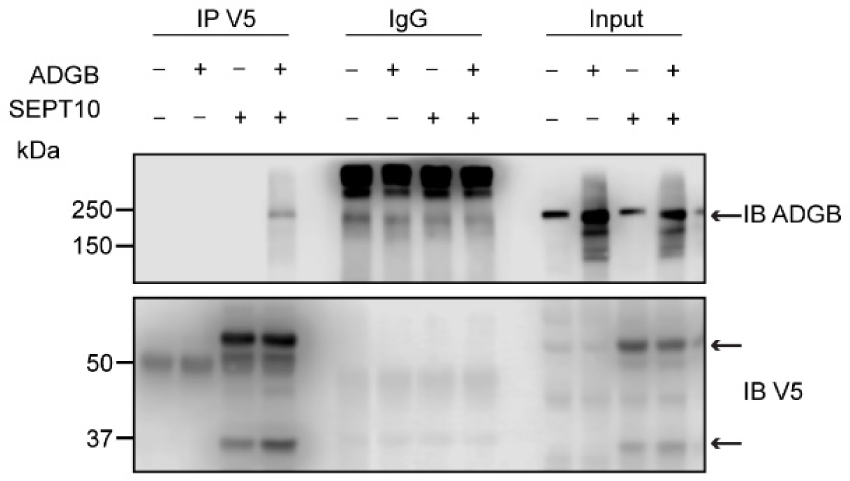
Reciprocal co-immunoprecipitation (coIP) of ADGB and V5-SEPT10 from. Fig 7B. Representative immunoblot of ADGB and V5 in protein lysates of HEK293 cells (co-)transfected with full-length ADGB and a C-terminally V5-tagged SEPT10 construct following co-immunoprecipitation of ADGB and V5-SEPT10.

**Figure S9.**
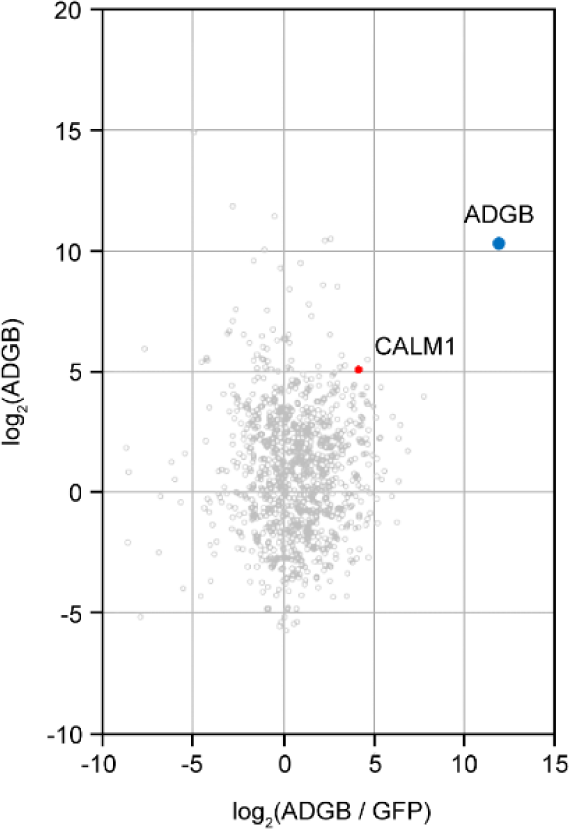
ADGB globin immunoprecipitation (IP) vs GFP control IP. Calmodulin (CALM1) is among the proteins that are prominently enriched in the anti-GFP IP of cells expressing the GFP-tagged globin domain of ADGB. The iBAQ values of the two IPs were log2 transformed and normalized against the median value. The normalized abundance of each protein detected in the globin IP (log2 globin) of cells expressing the GFP-globin construct is plotted against its specific enrichment compared to the control IP (log2 (globin / control)) from cells expressing the GFP-linker-GFP construct. ADGB and CALM1 are highlighted as blue and red dots, respectively, in the christmas tree plot representation.

**Figure S10.**
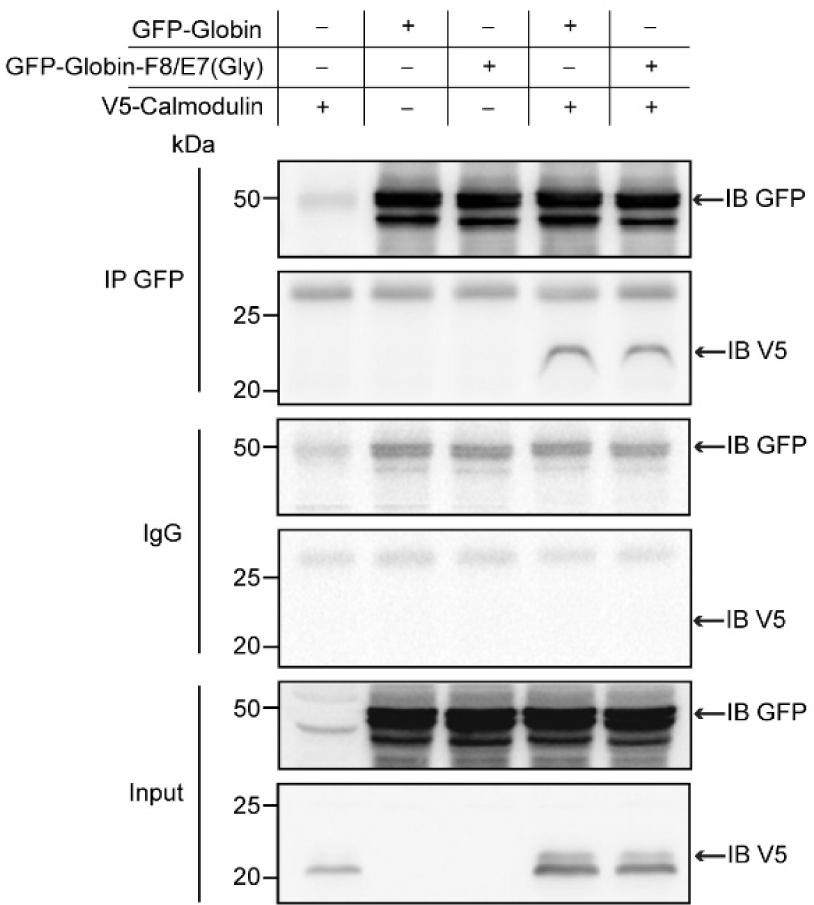
Double mutation of the key heme-binding residues does not abrogate interaction of ADGB and CaM. Representative immunoblot of GFP and V5 in protein lysates of HEK293 cells (co-)transfected with GFP-Globin, a GFP-Globin construct with a double mutation of the proximal heme-binding histidine and the distal heme-binding glutamine (GFP-Globin-F8/E7(Gly)) and V5-CaM following immunoprecipitation of GFP.

**Table S1.**
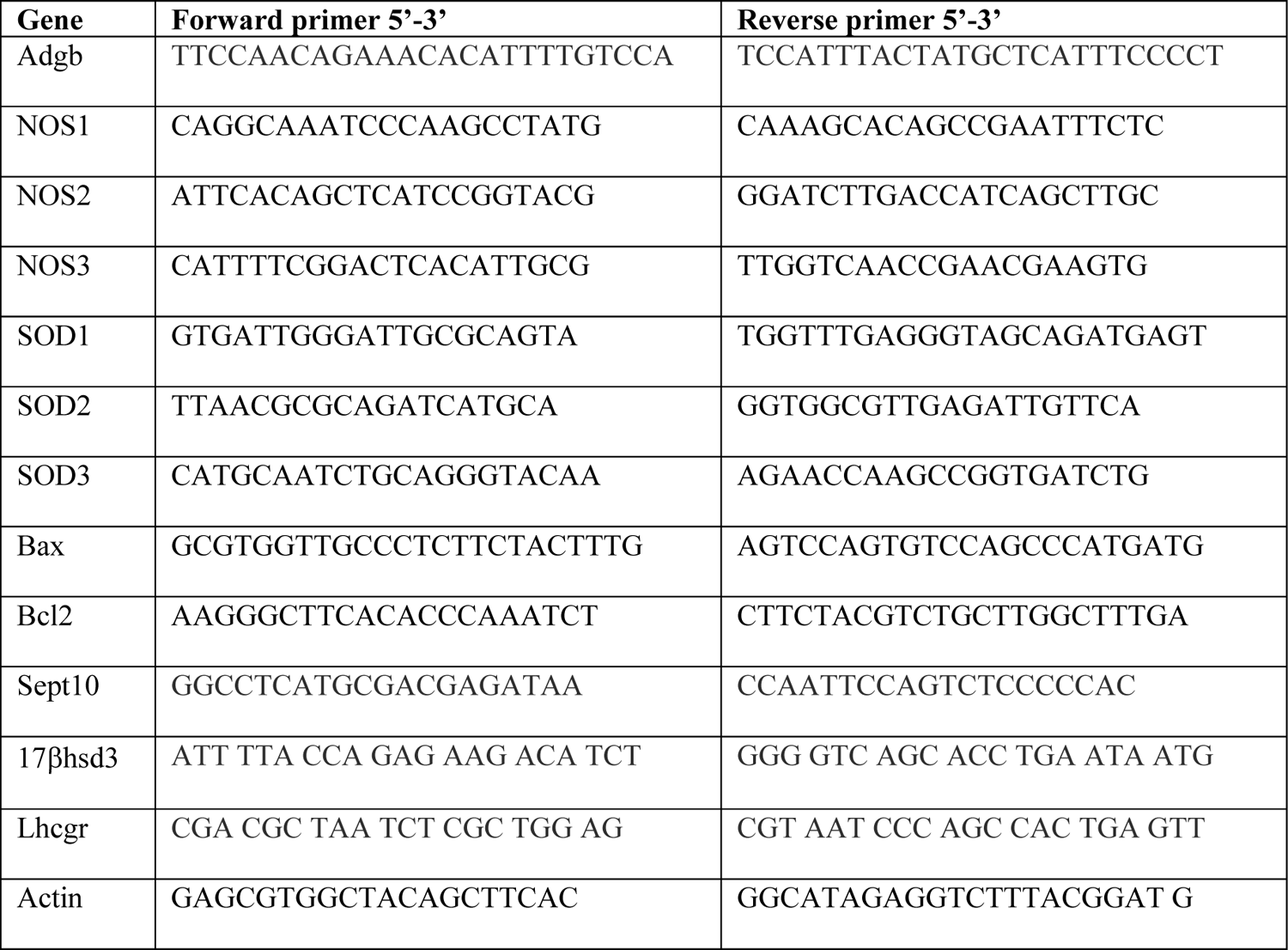
List of mouse primers used for RT-qPCR

**Table S2.** List of antibodies used throughout the study.

| **Antibody** | **Reference** | **Species** | **Application/dilution** |
| --- | --- | --- | --- |
| anti-mAdgb | Proteintech, custom made against region 409-745 of mAdgb | Rabbit | Immunoblot 1/200<br>IF 1/300 |
| anti-hAdgb N-ter | Atlas Antibodies, HPA036340 | Rabbit | Immunoblot 1/500 |
| anti-hAdgb C-ter | OriGene, TA330746 | Rabbit | Immunoblot 1/500 |
| anti-Sept10 | Proteintech, 12420-1-AP | Rabbit | Immunoblot 1/500<br>IF 1/300 |
| anti-Sept2 | Proteintech, 60075-1-Ig | Mouse | Immunoblot 1/500 |
| anti-Sept7 | Proteintech, 13818-1-AP | Rabbit | Immunoblot 1/500 |
| anti-Sept8 | Proteintech, 11769-1-AP | Rabbit | Immunoblot 1/500 |
| anti-Sept9 | Proteintech, 10769-1-AP | Rabbit | Immunoblot 1/500 |
| anti-Sept11 | Proteintech, 14672-1-AP | Rabbit | Immunoblot 1/500 |
| anti-Sept14 | Proteintech, 24590-1-AP | Rabbit | Immunoblot 1/500 |
| anti-GFP | Proteintech, 50430-2-AP | Rabbit | Immunoblot 1/1000 |
| anti-V5 | Invitrogen, 46-0705 | Mouse | Immunoblot 1/1000 |
| anti-CoxIV | Abcam, ab14744 | Rabbit | IF 1/300 |
| anti-HIF-1α | BD Transduction laboratories, 610958 | Mouse | Immunoblot 1/500 |
| anti-PHD2 | Novus Biologicals, NB100-137 | Rabbit | Immunoblot 1/1000 |
| anti-α-tubulin | Santa Cruz, TU-02 | Mouse | Immunoblot 1/1000 |

## Datasets

**Dataset S1.** Differentially regulated genes in wild-type versus Adgb knockout testes. Genes with a significant >2-fold induction or reduction are displayed.

**Dataset S2.** Raw MS data of the Adgb IP vs IgG control IP.

**Dataset S3.** Raw MS data of the ADGB globin IP vs GFP control IP.

## Source data files

**Source data files Figure 1.**
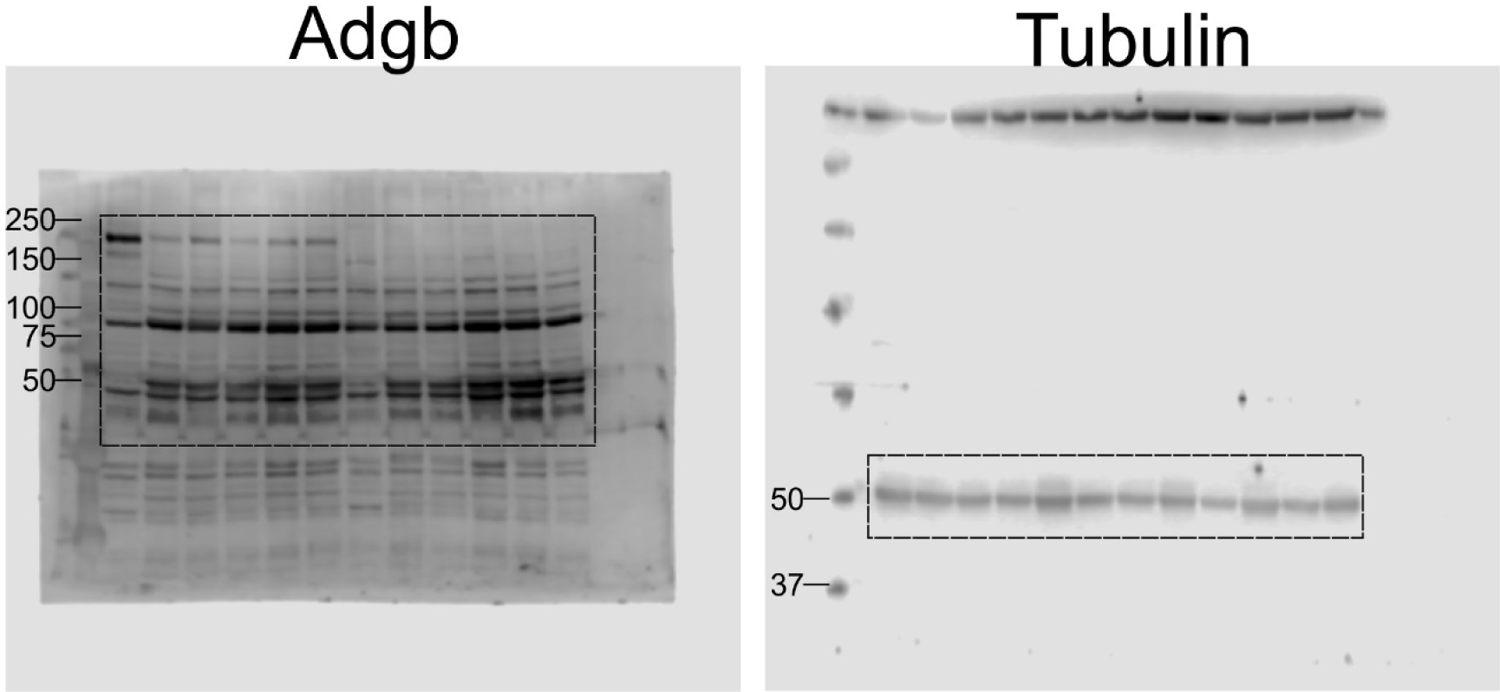
Full uncropped immunoblots for Adgb and tubulin.

**Source data files Figure 2.**
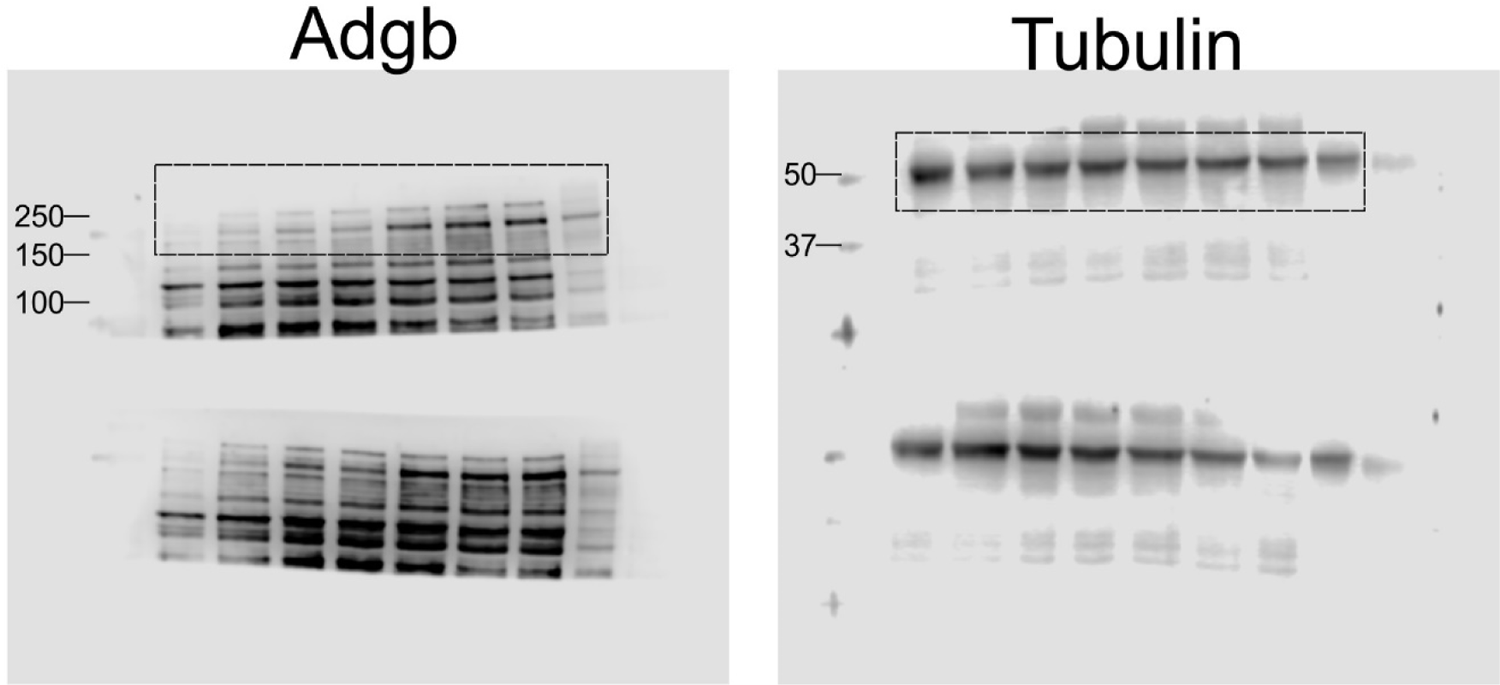
Full uncropped immunoblots for Adgb and tubulin.

**Source data files Figure 4B and 4C.**
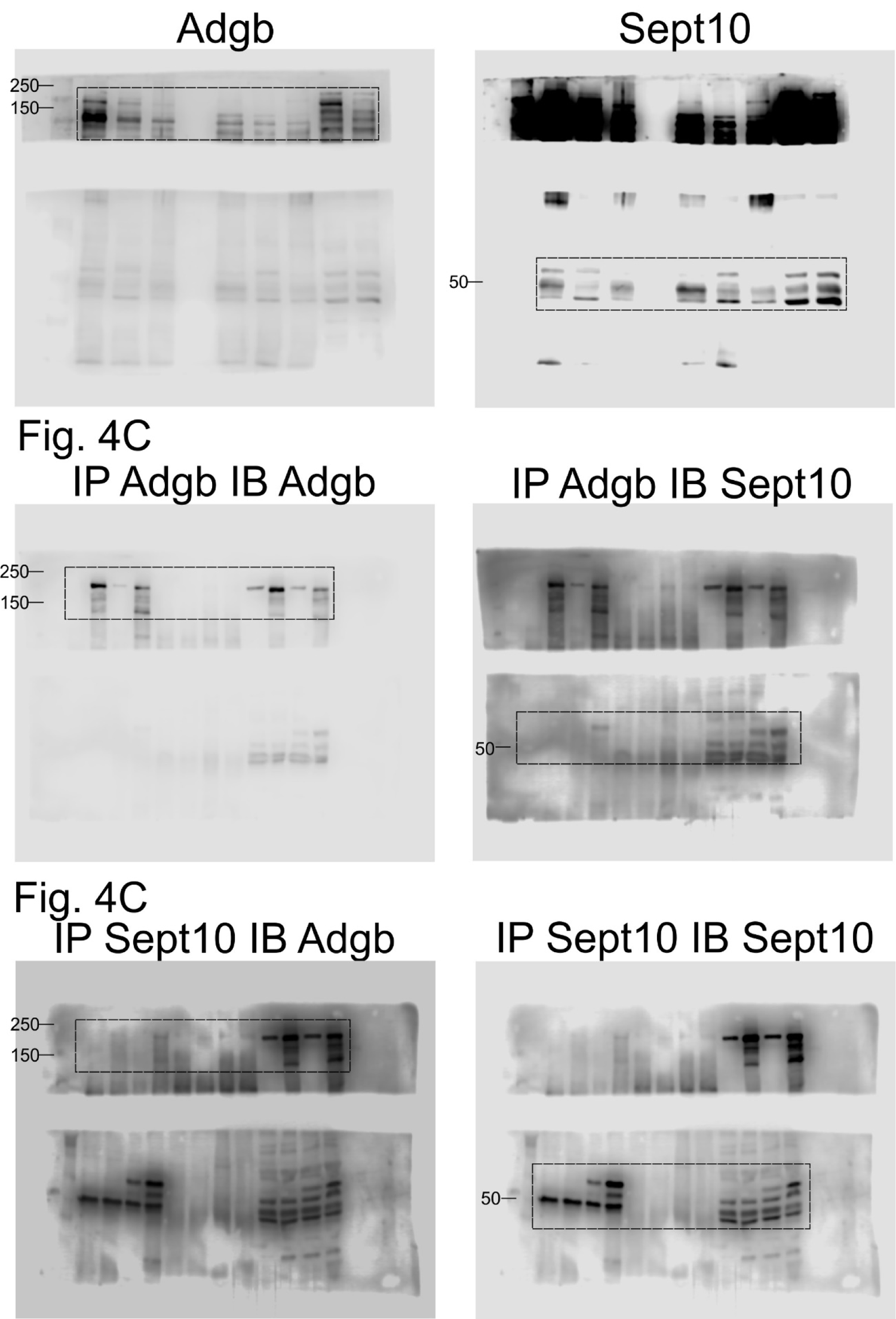
Full uncropped immunoblots for Adgb and Sept10.

**Source data files Figure 4D.**
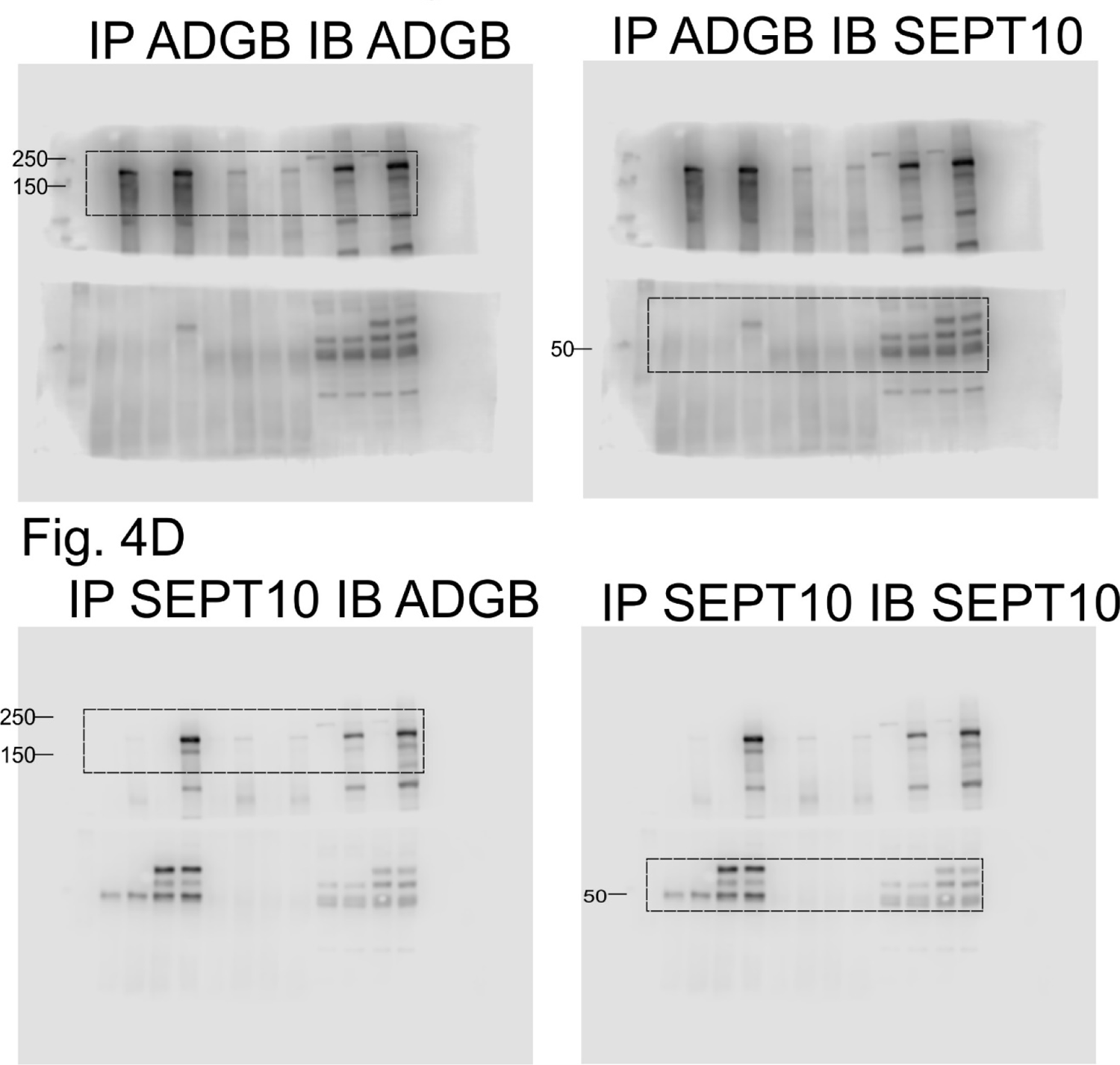
Full uncropped immunoblots for ADGB and SEPT10.

**Source data files Figure 4E.**
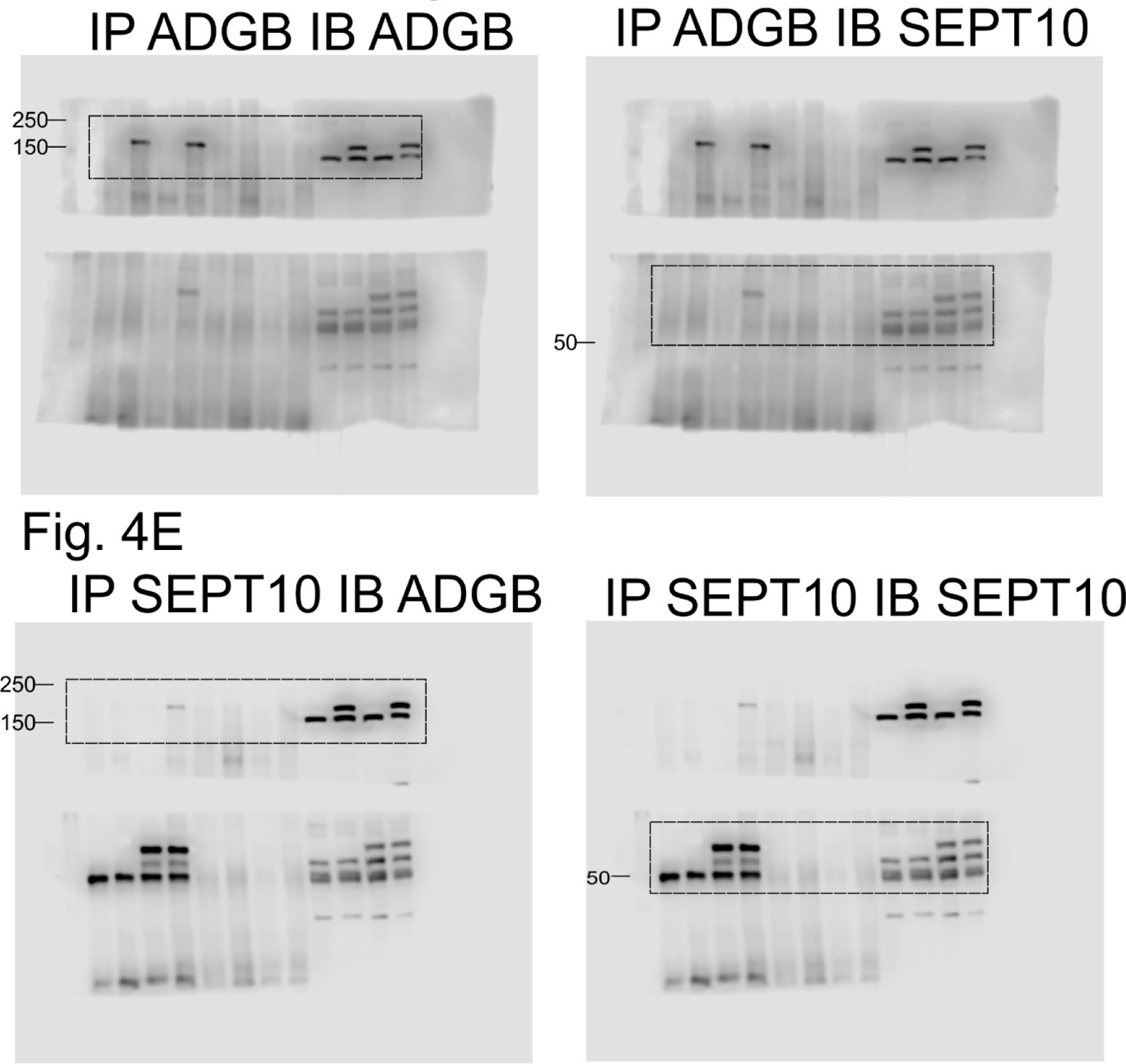
Full uncropped immunoblots for ADGB and SEPT10.

**Source data files Figure 6A and 6B.**
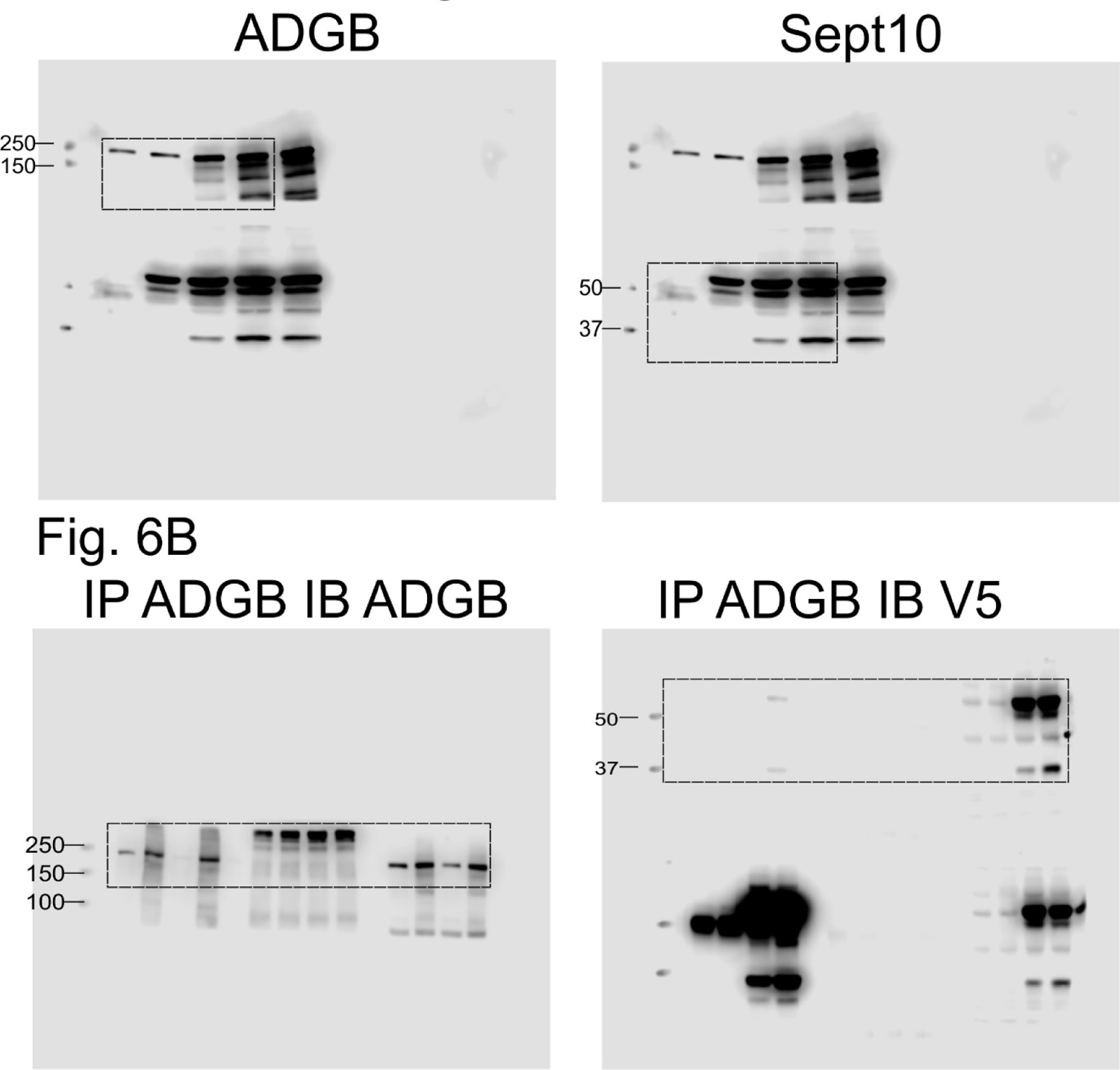
Full uncropped immunoblots for ADGB, V5 and SEPT10.

**Source data files Figure 6C and 6D.**
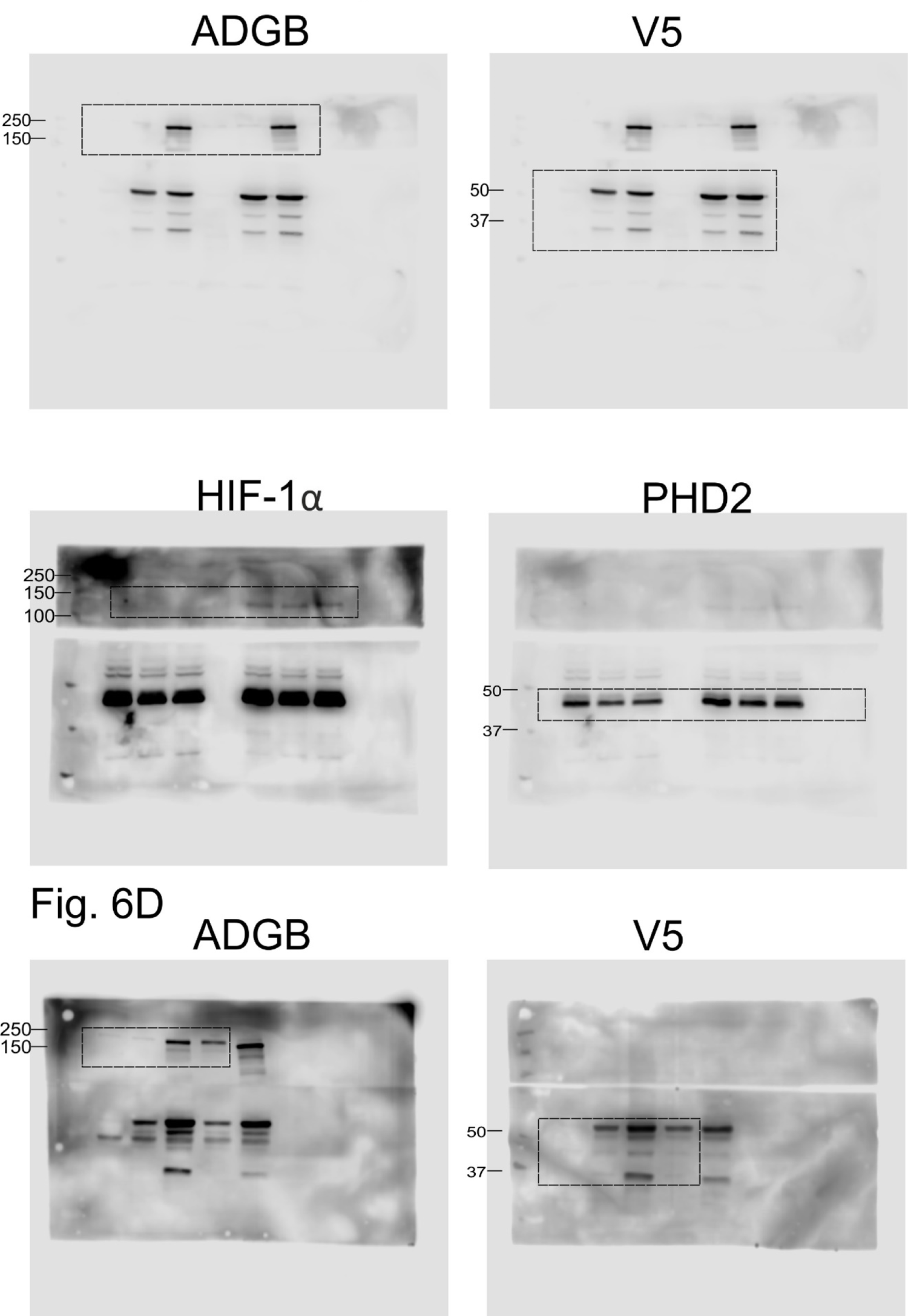
Full uncropped immunoblots for ADGB, PHD2, HIF-1α and V5.

**Source data files Figure 6E.**
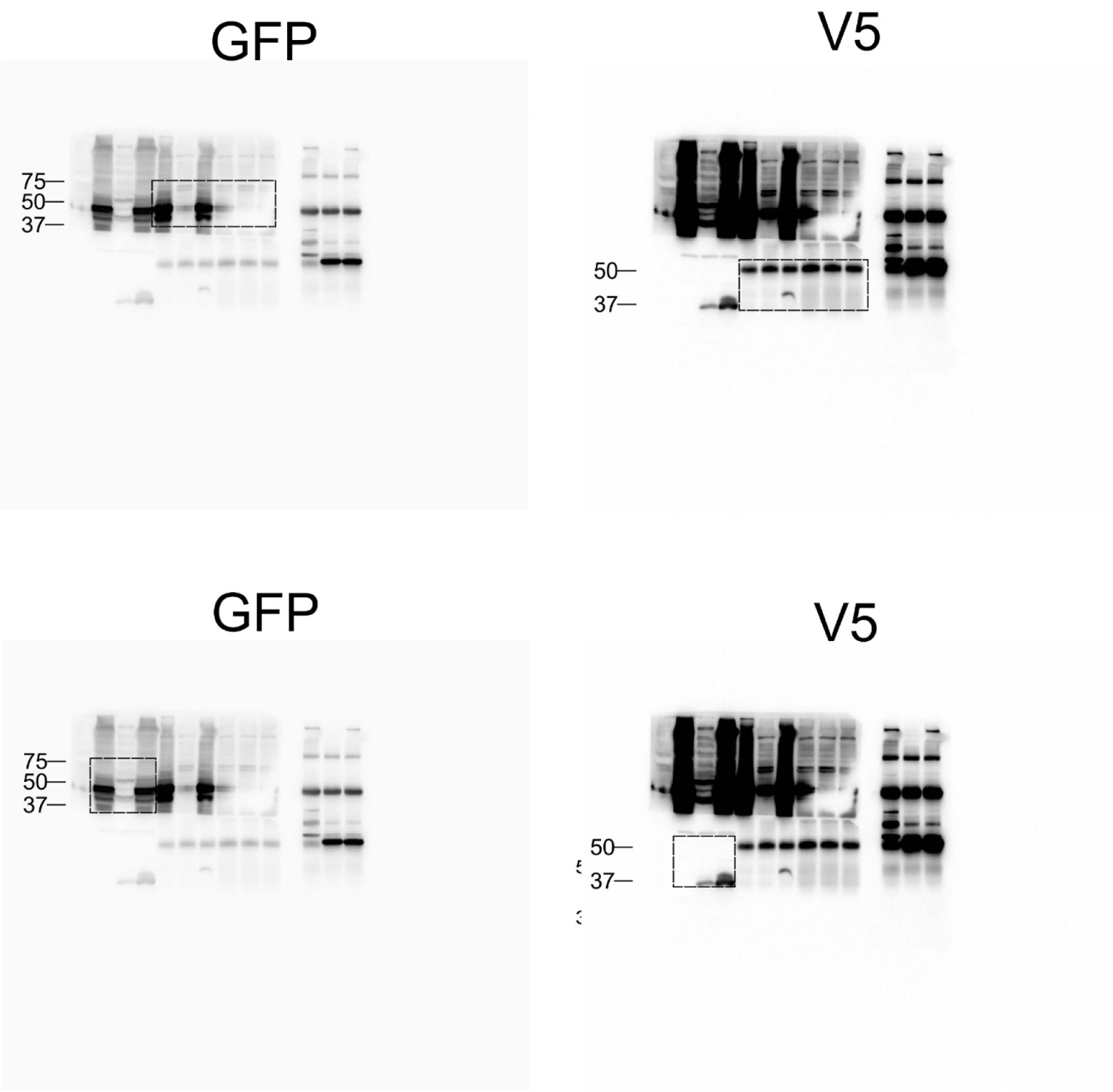
Full uncropped immunoblots for GFP and V5.

**Source data files Figure 6F.**
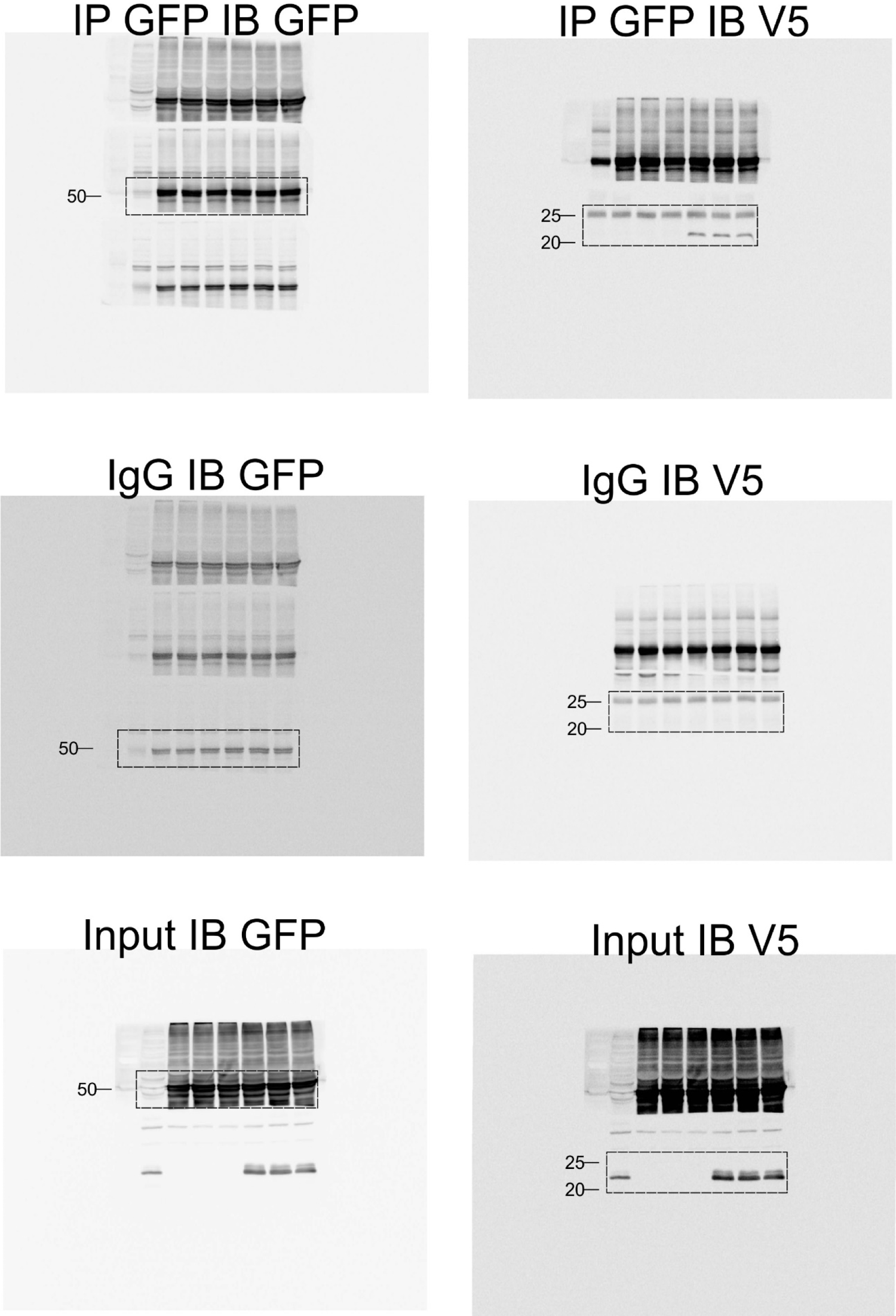
Full uncropped immunoblots for GFP and V5.

**Source data files Figure S1.**
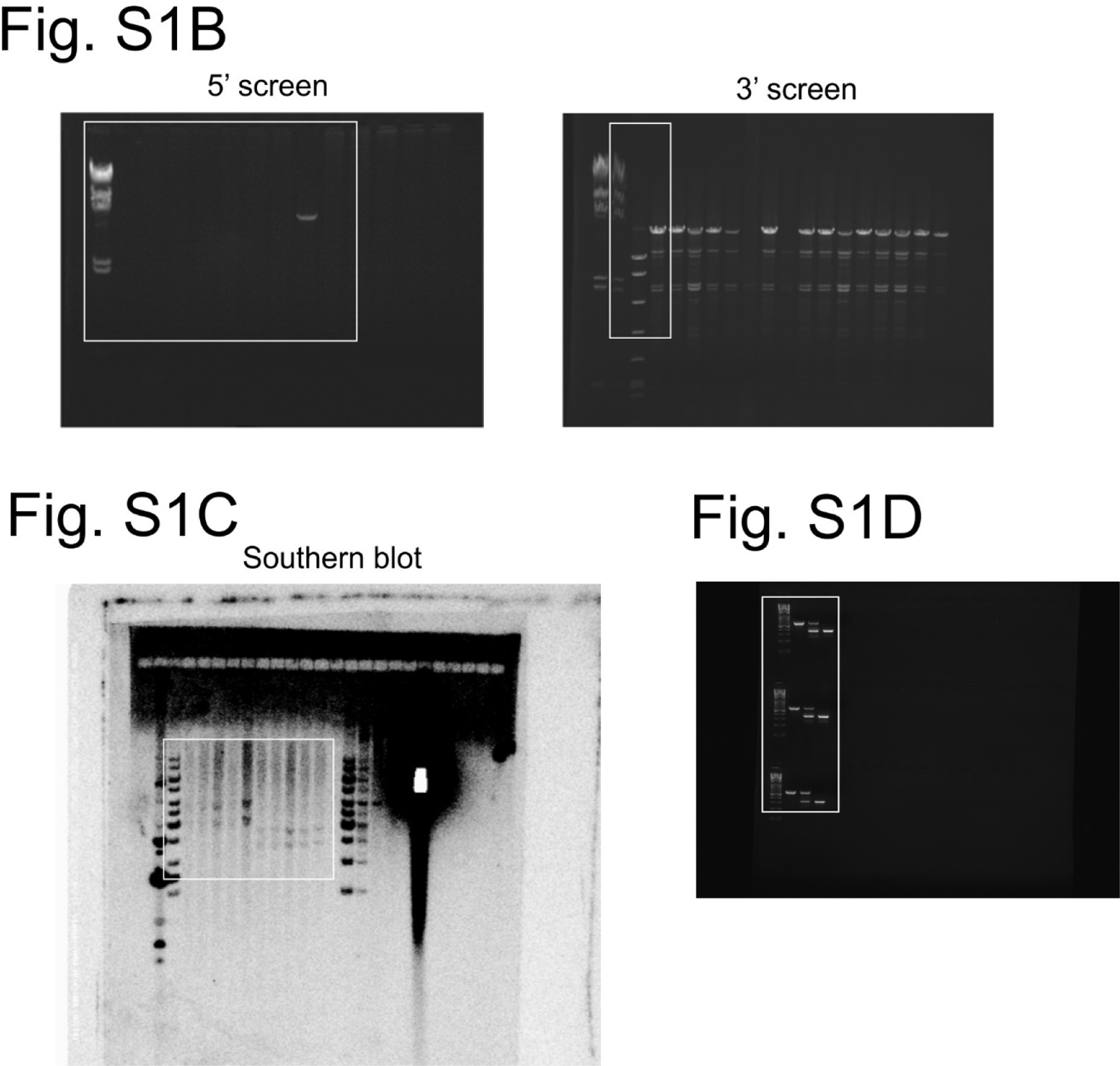
Full uncropped DNA gels and southern blot.

**Source data files Figure S5A, B, C.**
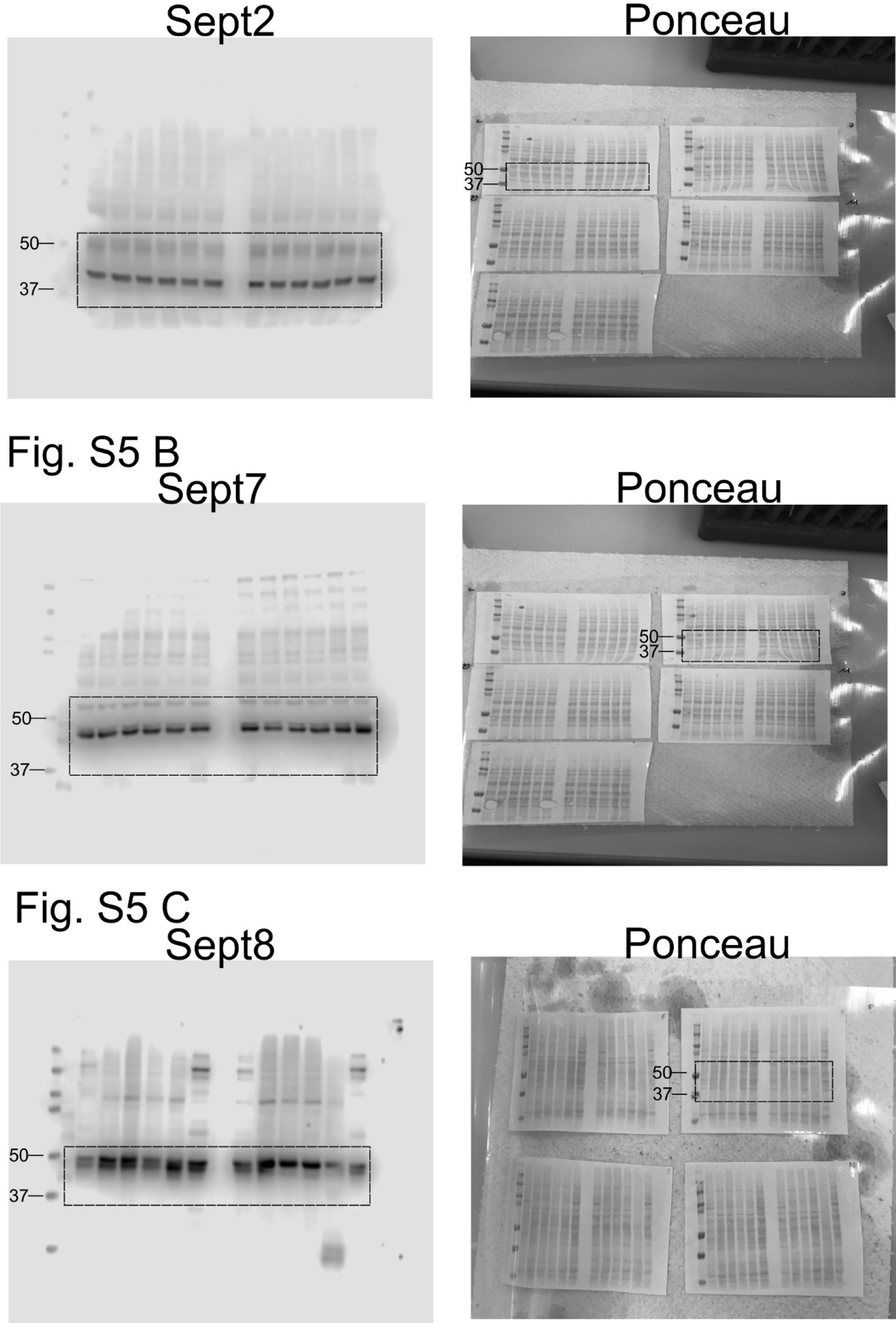
Full uncropped immunoblots for Sept2, Sept7, Sept8 and Ponceau S staining.

**Source data files Figure S5D, E, F.**
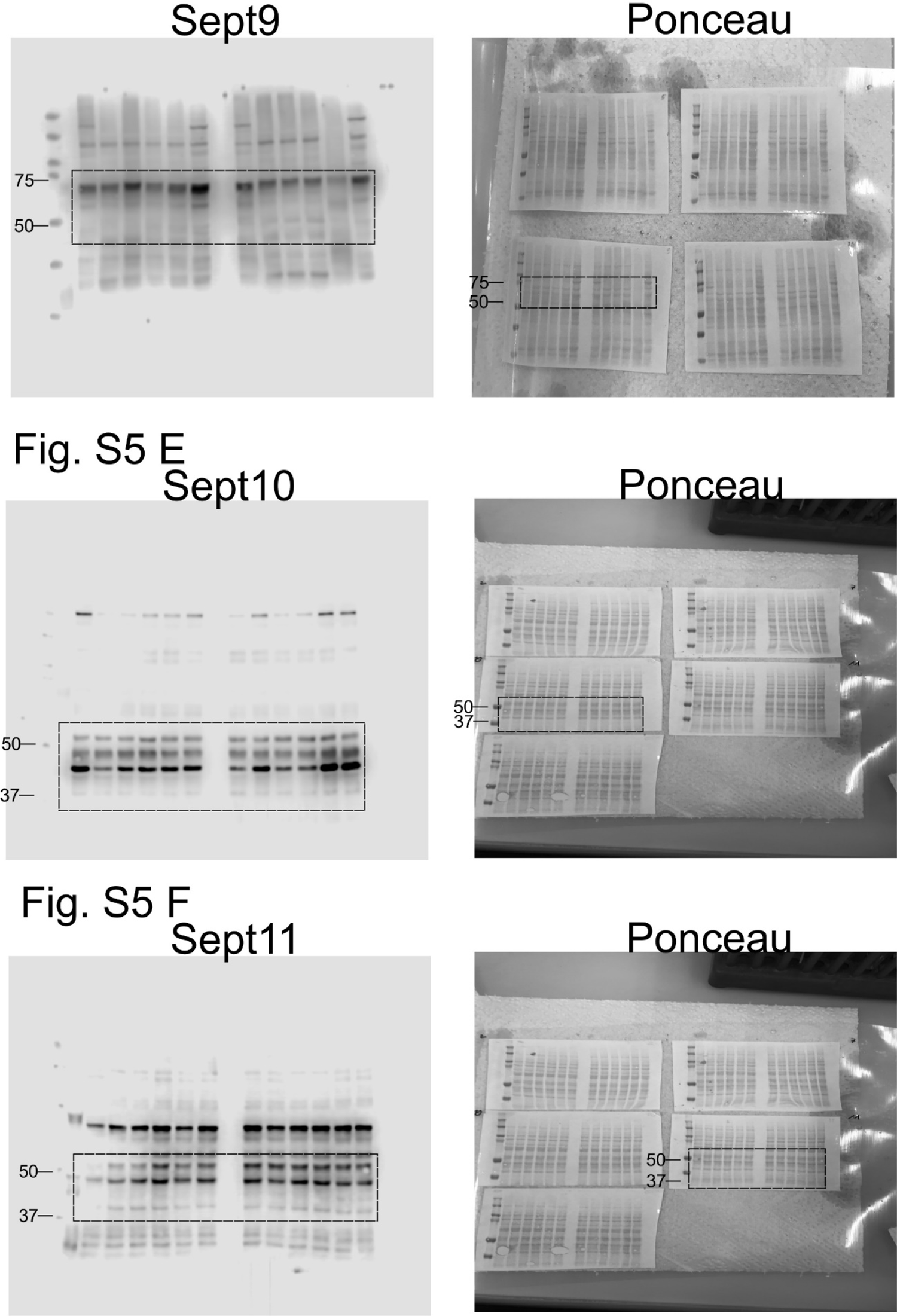
Full uncropped immunoblots for Sept9, Sept10, Sept11 and Ponceau S staining.

**Source data files Figure S5G.**
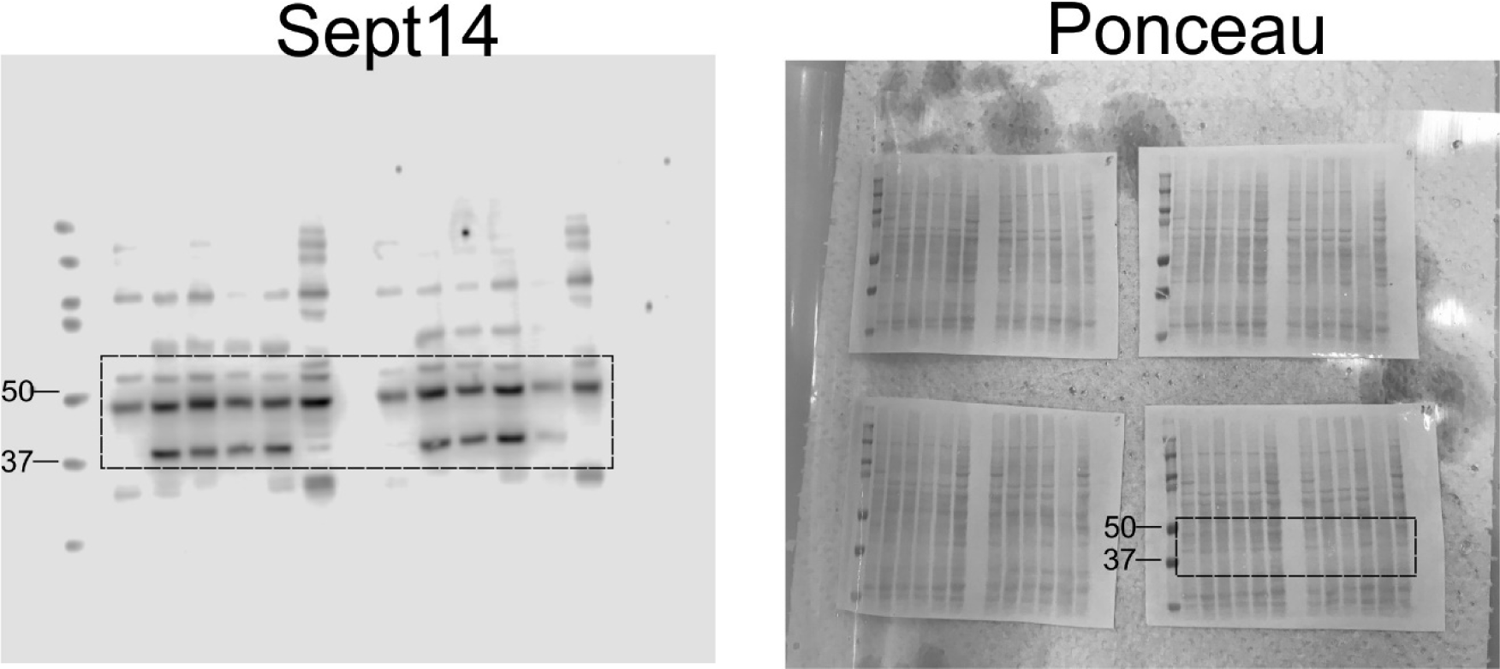
Full uncropped immunoblots for Sept14 and Ponceau S staining.

**Source data files Figure S6.**
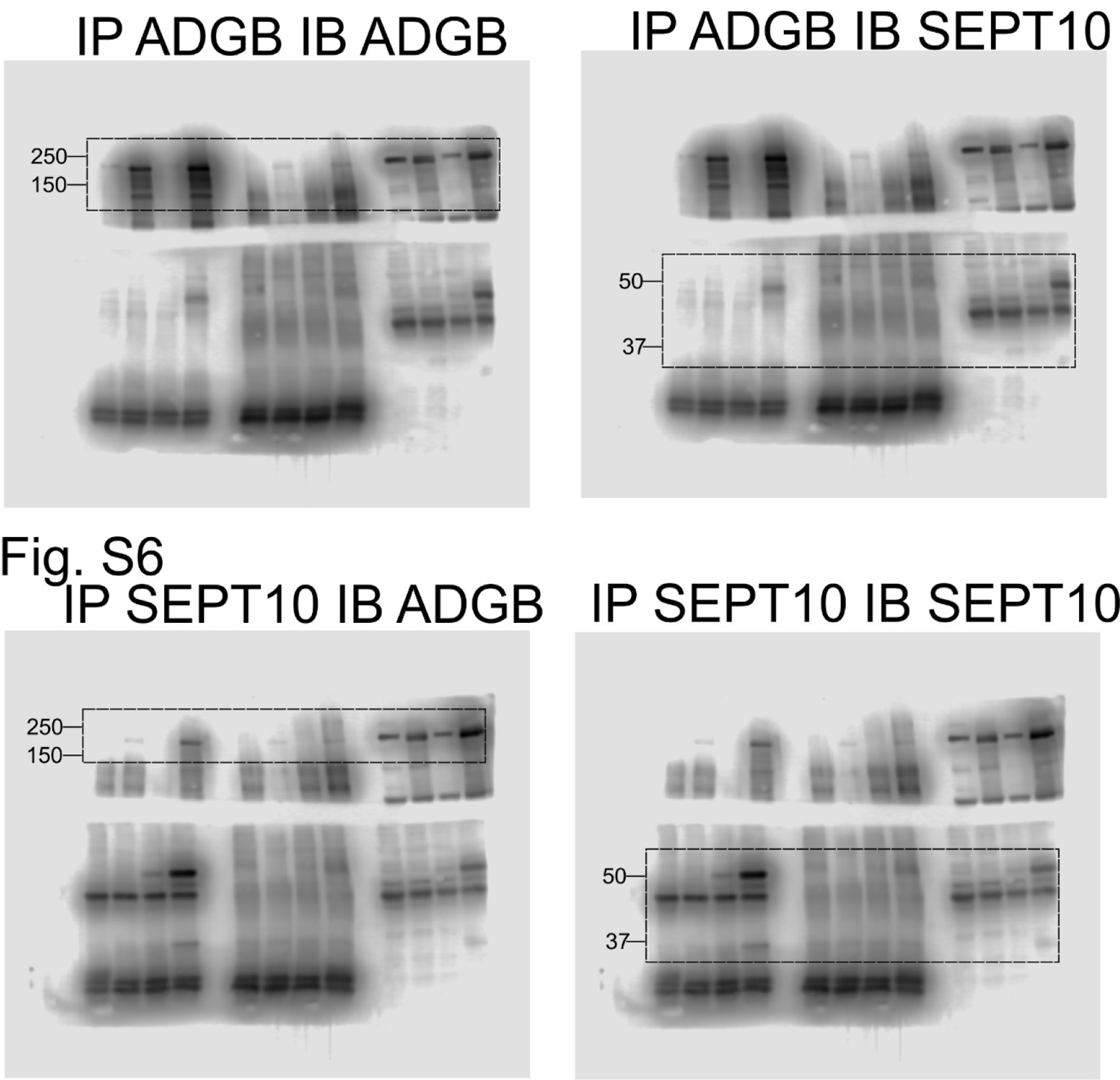
Full uncropped immunoblots for ADGB and SEPT10.

**Source data files Figure S7B.**
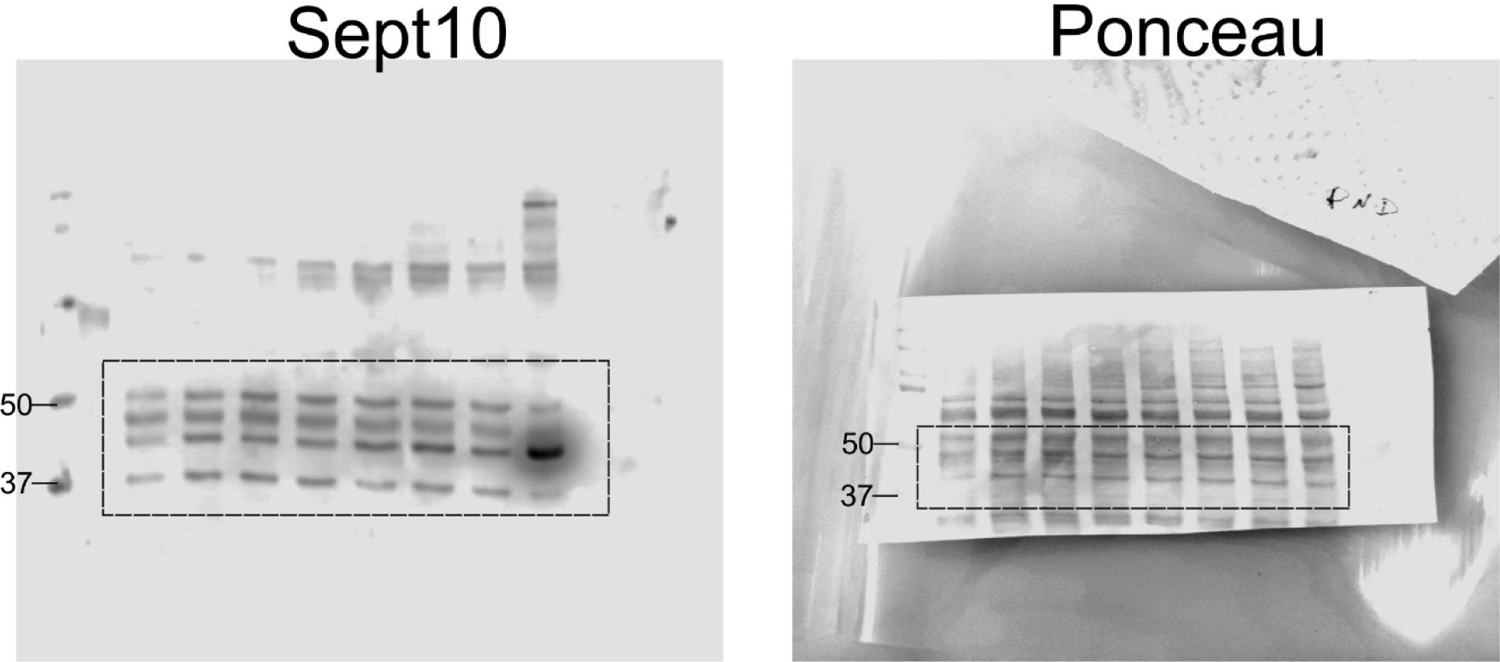
Full uncropped immunoblots for Sept10 and Ponceau S staining.

**Source data files Figure S8.**
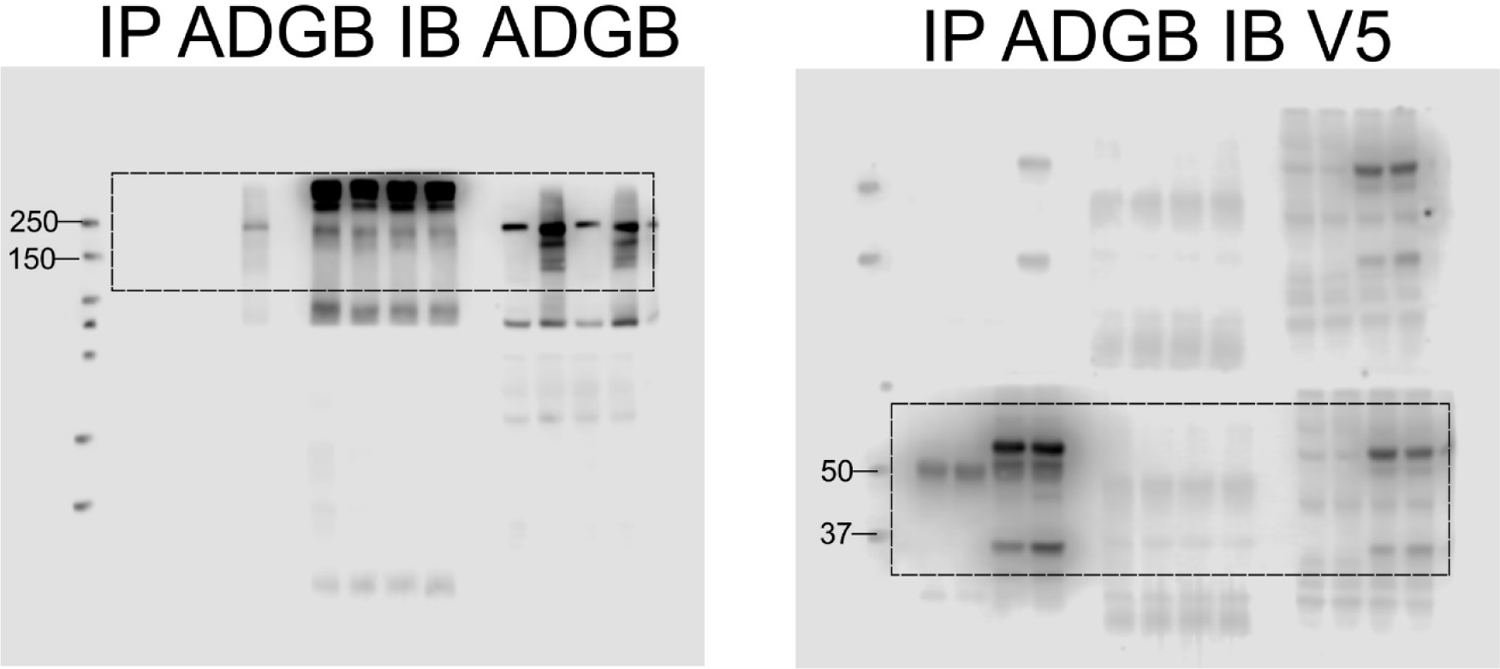
Full uncropped immunoblots for ADGB and V5.

**Source data files Figure S10.**
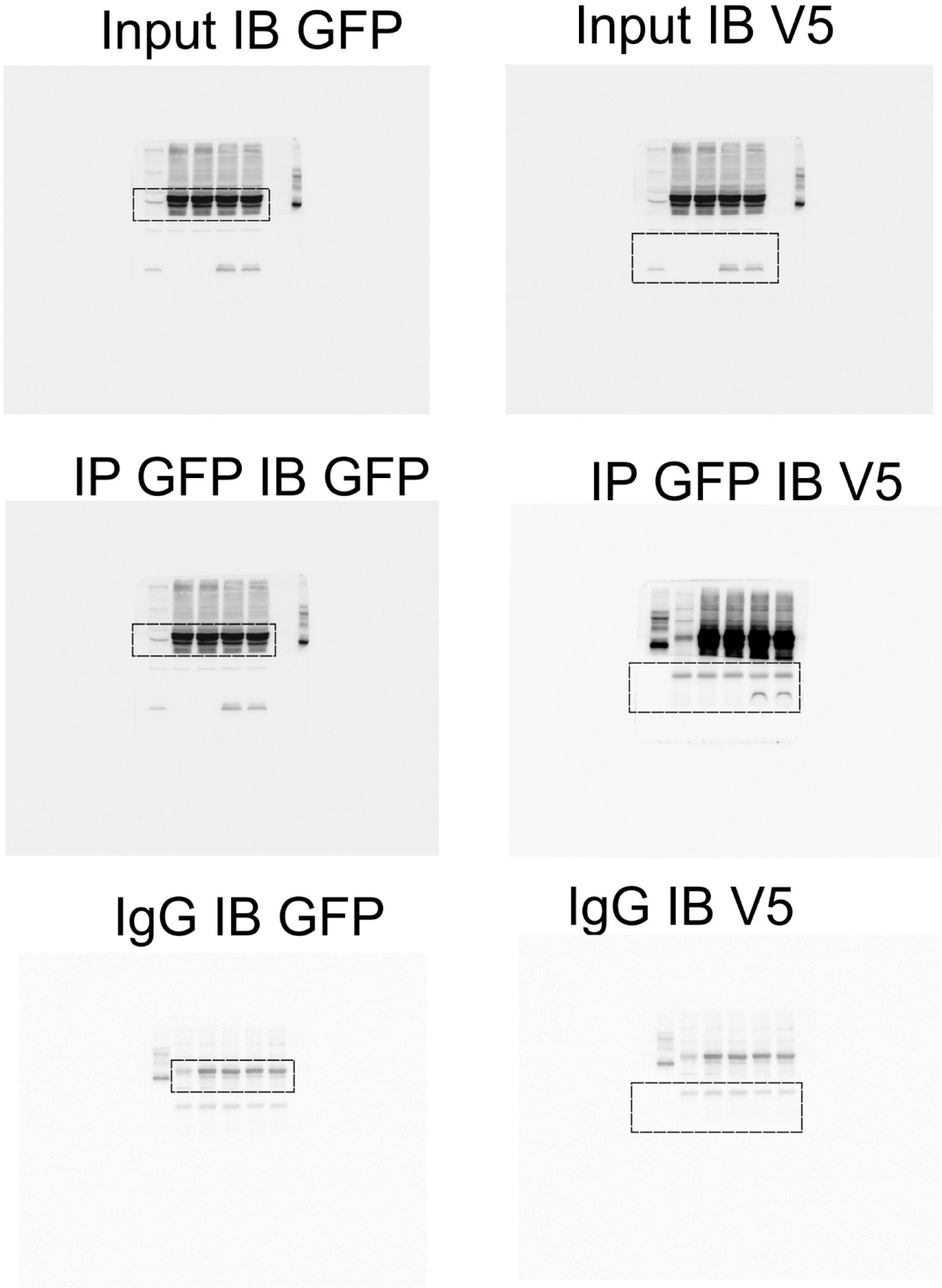
Full uncropped immunoblots for GFP and V5.

